# Structural Determinants of Synaptic Vesicle Protein 2C Ligand Selectivity and Their Impact on Dopamine Release

**DOI:** 10.64898/2026.09.10.750697

**Authors:** Matthew F. Martin, Meghan L. Bucher, Sejoon Choi, Anshumali Mittal, Eugene Mosharov, Adrian Hall, Christian Wolff, Michel Gillard, Laurent Provins, Peter S. Horanyi, David Sulzer, Gary W. Miller, Jonathan A. Coleman

## Abstract

The synaptic vesicle protein 2 (SV2) family is a highly conserved group of transmembrane glycoproteins sharing approximately 70% sequence identity. SV2A and SV2B are broadly distributed throughout the brain, while SV2C, the evolutionarily oldest member of the family, is primarily found in dopaminergic brain regions in the basal ganglia. Genome-wide association studies have linked SV2C to Parkinson’s disease, and SV2C appears to enhance dopamine storage in synaptic vesicles, but the basis for this effect is unknown. SV2s are the target of the racetam class of antiseizure medications, and SV2A-specific ligands are widely used to treat seizures. Recently several SV2C-specific ligands have been developed, but the structural basis for ligand specificity is unclear. A better understanding of SV2C structure, function, and pharmacology could lead to better targeted therapies for epilepsy, Parkinson’s disease, and other dopamine-related conditions. Here we present cryo-EM structures of apo SV2C, SV2C bound to the high affinity non-selective SV2 ligand padsevonil, SV2C bound to the SV2C-selective ligand UCB-F, and SV2A bound to the SV2A-selective ligand plosaracetam (also known as ABBV-552/SDI-118). We find that SV2C has a wider luminal opening than SV2A and SV2B, which allows for UCB-F to bind to the primary site and form favorable interactions that are not possible in the narrower primary binding site of SV2A and SV2B. We also demonstrate that UCB-F and padsevonil, but not plosaracetam, reduce dopamine release in striatal sections of mouse brain. Our biochemical experiments and structures provide insights into SV2 ligand specificity and offer a template for the rational development of therapeutics targeting SV2C.

## Introduction

The synaptic vesicle protein 2 (SV2) family is a group of transmembrane glycoproteins – SV2A, SV2B, and SV2C. SV2A, the most abundant isoform, is ubiquitous in the adult brain and is also found in endocrine tissues such as the pancreas and adrenal medulla^1–3^. SV2B is found in most brain regions, and is enriched in the retina, while SV2C is the least abundant member of the family and is primarily expressed in dopaminergic regions in the basal ganglia^1,4–8^. SV2C-like genes can be found in invertebrates such as *Drosophila*, and all three SV2 isoforms first appeared in cartilaginous fish^9^. The high degree of SV2 conservation in all organisms from fish to mammals suggests that these proteins carry out an important function that likely emerged concurrently with, or shortly after, the evolution of vertebrates.

Since the initial discovery and cloning of the SV2A gene in the 1990s, the function of the SV2 family has been a matter of intense speculation^9–11^. SV2s share significant sequence homology with major facilitator superfamily transport proteins, and were first thought to be neurotransmitter transporters given their location in synaptic vesicles^2,10,12^. Knockout of SV2A in mice results in a lethal epilepsy phenotype, and humans with SV2A mutations suffer from intractable seizures and developmental disabilities, supporting the hypothesis that SV2s have a critical function in the brain^13–16^. Although several putative SV2 substrates have been suggested over the years, no transport substrates have been conclusively identified for any SV2 family member to date, and the function of SV2s remains unclear^17,18^. All three SV2 family members bind to and traffic synaptotagmins (Syts), crucial regulators of vesicle exocytosis, but antiseizure medications do not appear to modulate these interactions, and mutation of conserved transmembrane residues hypothesized to be involved in substrate binding and transport causes aberrant neurotransmission without affecting Syt trafficking, suggesting that SV2s have multiple functions in neurons^19–25^. SV2s also serve as the entry receptors for the bacterial neurotoxins tetanus toxin (TeNT) and botulinum toxin (BoNT, clinically used as Botox), which bind to the glycosylated luminal domain in order to enter neurons via endocytosis^26–31^. SV2A is the target of the widely used antiseizure medications (ASMs) levetiracetam (LEV,) and brivaracetam (BRV,), and pan-SV2 ligands such as padsevonil (PSL) have been tested clinically for treatment of focal seizures^32–39^. The SV2A-selective ligand plosaracetam (PRM, also known as ABBV-552/SDI-118) is in clinical trials for use as a cognitive enhancer to treat symptoms of Alzheimer’s disease and dementia^40,41^. Recently, several structures of SV2A and SV2B in complex with these ligands have been published^30,42–46^. These structures reveal that drugs bind to a primary site located in the transmembrane domain, consistent with the hypothesis that they act as transport inhibitors^42–46^. The primary drug binding site is remarkably conserved amongst SV2s, and due to the lack of structural information for SV2C, the basis for the wide range of racetam affinity and selectivity across the SV2 family remains unclear.

Although the precise function of SV2s is still unknown, recent studies have linked SV2C to the pathogenesis and treatment of Parkinson’s disease (PD), a neurological disorder caused by loss of dopaminergic neurons in the substantia nigra which results in movement disorders^47–49^. It has been suggested that PD is related to the disruption of proper dopamine storage in synaptic vesicles, leading to neuronal cell damage and death caused by oxidative stress^50^. Variation in the SV2C gene have been suggested to predict the efficacy of L-DOPA treatment in PD patients and treatment with antipsychotics in patients with schizophrenia^51,52^. SV2C variants are also reported to influence the occurrence of adverse events linked to dopamine therapies^53^. Studies in *Drosophila* indicate that nicotine, which has long been recognized for its protective effect against PD, requires expression of the SV2C orthologues SV2L1 and SV2L2 to rescue flies from a model PD phenotype caused by overexpression of alpha-synuclein, and to protect against paraquat-induced PD^54–56^. Co-expression of human SV2C with the vesicular monoamine transporter 2 (VMAT2), which is responsible for loading of dopamine into synaptic vesicles, enhances neurotransmitter loading *in-vitro*^47^.

Postmortem analysis of PD patient brain tissue shows disruptions in SV2C expression, and genetic deletion of SV2C in mice results in impaired dopamine release^48^. These observations suggest that SV2C plays an important, yet unknown function in dopamine homeostasis, vesicle loading, and release, and that pharmacological control of this function could serve as a novel PD therapy. Only one SV2C-specific ligand, UCB-F, has been developed, and due to the high degree of conservation between the primary binding site and the lack of published SV2C structures, the structural basis for UCB-F selectivity remains unknown^57^. SV2A-selective ligands have also garnered interest as cognitive enhancers. The recently developed SV2A-specific ligand plosaracetam (PRM), which is currently in clinical trials for treatment of Alzheimer’s disease and dementia symptoms, is reported to be a cognitive enhancer with no antiseizure effects, despite sharing many features with antiseizure racetam drugs^40,41^. It is not known why PRM, but not other ligands, lacks antiseizure activity, and like UCB-F, little information is known about its selectivity and kinetics. Furthering our understanding of SV2 pharmacology and function will allow the development of new targeted therapies for epilepsy, PD, and other neurological conditions, and will provide new insights into the pathology of these diseases as well as neurotransmission in the healthy brain.

Here we present three single particle cryo-electron microscopy (cryo-EM) structures of SV2C, in apo-, PSL-bound, and UCB-F-bound states, along with a structure of SV2A bound to the SV2A-selective cognitive enhancer molecule PRM. These structures reveal structural features that underpin ligand specificity for the SV2 family and allow for detailed comparisons amongst SV2 family members. Biochemical experiments using radiolabeled PSL reveal that non-conserved residues in the primary binding site of SV2C induce more favorable binding site occlusion in the UCB-F-bound state. Fast scan cyclic voltammetry studies demonstrate that both UCB-F and PSL inhibit dopamine release in mouse brain slices, while studies in SV2C-expressing cells suggest that these ligands do not alter vesicular dopamine content, indicating potential compound-mediated effects by a different mechanism such as vesicular trafficking during neurotransmission.

## Results

### Characterization of SV2 ligands

Recently published structures of SV2A and SV2B revealed that the primary binding sites are nearly 100% conserved^30,42–46^. Sequence alignments show that SV2C is also similarly conserved. Despite the high degree of similarity between these proteins, there is a remarkable variation in affinity and specificity of racetam-type SV2 ligands, which all share a substituted 2-pyrrolidone core structure **(Figure 1A)**. The ASMs levetiracetam (LEV) and brivaracetam (BRV) are SV2A specific and are being used to treat seizures. LEV binds to SV2A with a K_d_ of approximately 2 µM, with negligible binding to SV2B and SV2C, and BRV binds to SV2A with a K_d_ of approximately 250 nM and SV2B with a K_i_ of approximately 70 µM^36–39,42,43^. Padsevonil (PSL), is non-selective high-affinity ligand and binds all three SV2s with nanomolar affinity **(Figure 1B)** and UCB-J, used as a PET tracer, binds SV2A with a K_d_ of approximately 1 nM, SV2B with a K_i_ of approximately 1 µM, and SV2C with a K_i_ of 750 nM **(Extended Data 1A**)^32–34,42,58,59^. Plosaracetam (PRM), evaluated in clinical trials as a cognitive enhancer for Alzheimer’s disease and dementia patients, is thought to be selective for SV2A, but no published studies to date have measured PRM binding to SV2s^40,41^. Although there are many SV2A-specific racetam ligands, only one SV2C-specific ligand, UCB-F, has been described in the literature, and there is also no published information available regarding its selectivity and binding characteristics^57^. To characterize these ligands in detail, we first performed competition binding experiments using SV2-enriched membranes and ^3^H-labelled PSL to determine the K_i_ of UCB-F and PRM for SV2A, SV2B, and SV2C. UCB-F binds to SV2C with a Ki of 37 nM, achieving approximately 20-fold selectivity over SV2A (755 nM), and binds very weakly to SV2B, as we were unable to reliably determine a K_i_ **(Figure 1C)**. PRM bound SV2A with a K_i_ of 5 nM (Kd 14 nM), demonstrating approximately 2000-fold selectivity over SV2B (12.8 µM) and approximately 1000-fold selectivity over SV2C (4.9 uM) **(Figure 1D)**.

**Figure 1.**
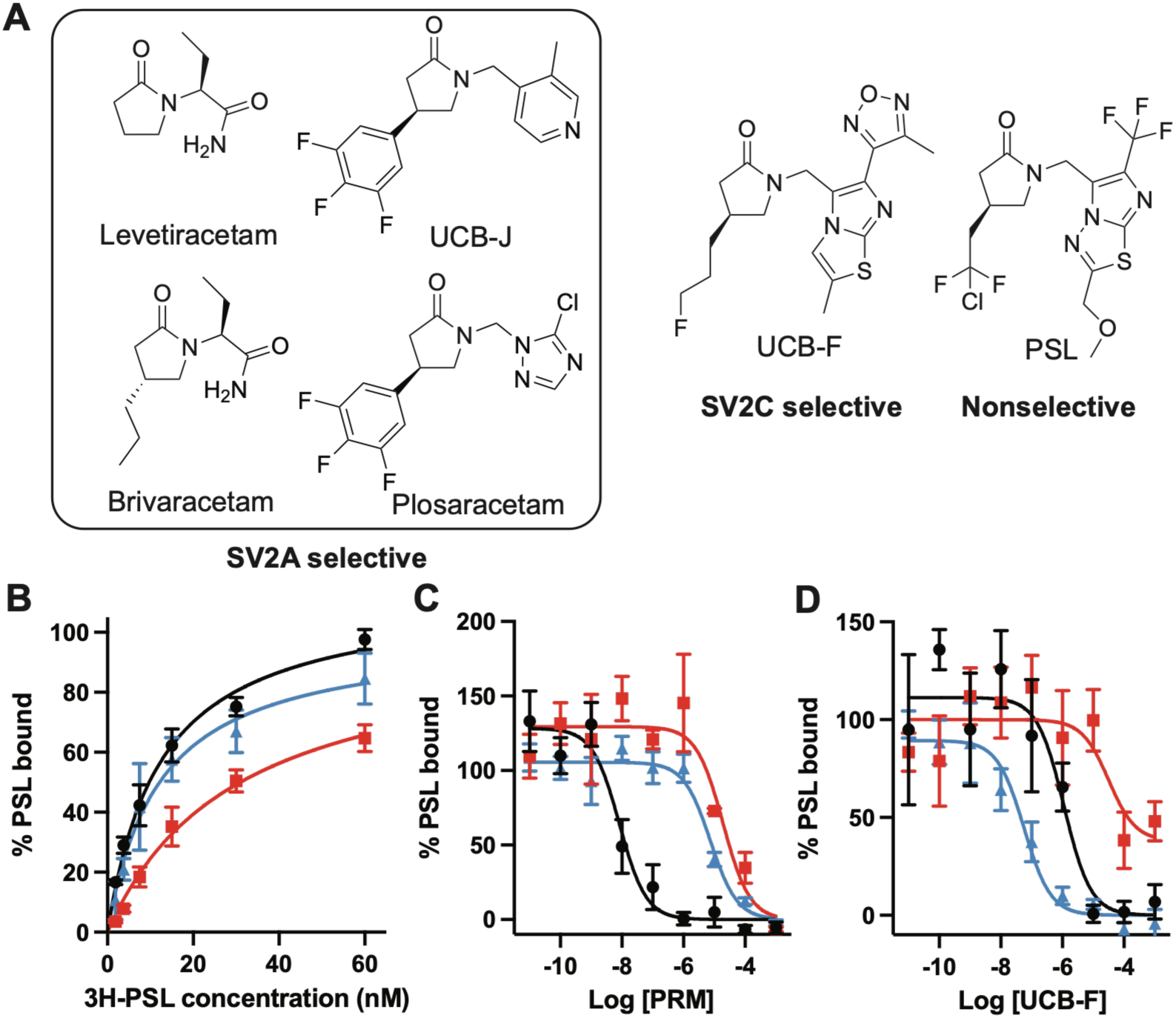
**A)** Structures of racetam ligands with varied selectivities for the SV2 family. **B)** Saturation binding of 3H-padsevonil to SV2A (black circles, K_d_ = 35 nM +/- 3 nM), SV2B (red squares, K_d_ = 31 nM +/- 3 nM), and SV2C (blue triangles, K_d_ = 12 nM +/- 2 nM). **C)** Competition binding of PRM with 5 nM 3H-padsevonil to SV2A (black circles, K_i_ = 5.5 nM +/- 1.4 nM), SV2B (red squares, K_i_ = 12.8 uM +/- 5.1 uM), and SV2C (blue triangles, K_i_ = 4.9 uM +/- 1.1 uM). **D)** Competition binding of UCB-F with 5 nM 3H-padsevonil to SV2A (black circles, K_i_ = 755 nM +/- 351 nM), SV2B (red squares, K_i_ not measurable), and SV2C (blue triangles, K_i_ = 37 nM +/- 10 nM).

Previous studies have shown that SV2C enhances VMAT2-mediated neurotransmitter loading in cultured cells^60^. To understand how ligand binding to SV2C affects this process, we performed experiments examining the effect of PSL, UCB-F, and PRM on the uptake of a fluorescent dopamine-like tracer known as false fluorescent neurotransmitter 206 (FFN206), which is accumulated by VMATs^61^. We generated cell lines that stably expressed either VMAT2 alone, or both VMAT2 and SV2C, and examined their ability to load FFN206 into vesicles. Co-expression of SV2C with VMAT2 increased FFN206 uptake by approximately 14% relative to cells expressing VMAT2 alone **(Figure 2A)**. When cells treated with high concentrations of PSL (50 µM PSL) we observed a 21% decrease in FFN206 loading relative to DMSO-treated cells, but this effect was only observed at the highest concentration of PSL which is in large excess of the K_i_ and not at lower concentrations **(Figure 2B)**. In support of these ligands having only minor effects on FFN206 loading, we found that UCB-F treatment did not affect FFN206 uptake **(Figure 2C**) and treatment of cells with PRM, an SV2A-specific ligand also had no effect **(Figure 2D**). Treatment of cells with PSL, UCB-F, and PRM also caused an increase in SV2C expression relative to DMSO treated HEK293 cells which was dose-dependent , suggesting that these ligands may have a stabilizing effect on SV2C **(Extended Data 2).**

**Figure 2.**
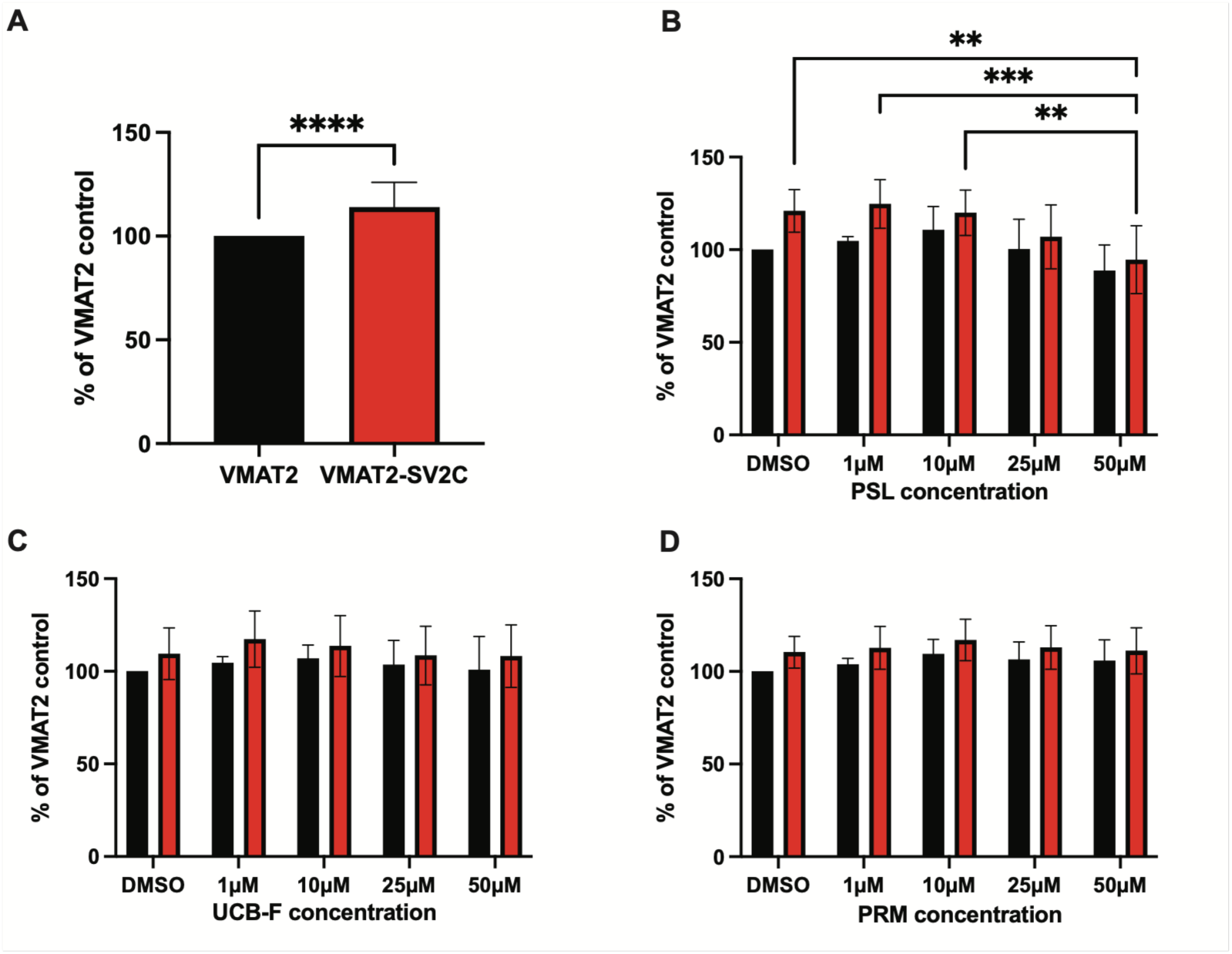
**A)** Effect of SV2C coexpression on VMAT2-mediated FFN-206 uptake. FFN206 uptake was normalized to 100% in HEK293 cells expressing VMAT2 (black bar). Coexpression with SV2C increased FFN206 uptake by 14% (red bar, p < 0.0001, Welch’s t test). **B)** Effect of PSL treatment on FFN206 uptake in VMAT2 control (black bars) and VMAT2-SV2C (red bars) cells. FFN206 uptake was normalized to 100% in HEK293 cells expressing VMAT2 and treated with DMSO. Treatment with 50 uM PSL caused a decrease in FFN206 loading by 21% relative to untreated cells. Significance was determined by two-way ANOVA with Šídák’s multiple comparisons test. Effect of cell line (F[1, 50] = 11.08, p = .0016); effect of treatment (F[4, 50] = 5.474, p = .0010); effect of interaction (F[4, 50] = 0.7526, p = .5610), n = 4 – 8 experimental replicates, **p < .01, ***p < .001). **C)** Effect of UCB-F treatment on FFN206 uptake in VMAT2 control (black bars) and VMAT2-SV2C (red bars) cells. FFN206 uptake was normalized to 100% in HEK293 cells expressing VMAT2 and treated with DMSO. No significant changes in FFN206 loading were observed. Significance was determined by two-way ANOVA with Šídák’s multiple comparisons test Two-way ANOVA with Šídák’s multiple comparisons test. Effect of cell line (F[1, 35] = 3.290, p = .0783); effect of treatment (F[4, 35] = 0.3686, p = .8294); effect of interaction (F[4, 35] = 0.08755, p = .9857), n = 3 – 6 experimental replicates, n.s.). **D)** Effect of PRM treatment on FFN206 uptake in VMAT2 control (black bars) and VMAT2-SV2C (red bars) cells. FFN206 uptake was normalized to 100% in HEK293 cells expressing VMAT2 and treated with DMSO. No significant changes in FFN206 loading were observed. Significance was determined by two-way ANOVA with Šídák’s multiple comparisons test. Effect of cell line (F[1, 50] = 7.603, p = .0081); effect of treatment (F[4, 50] = 0.8518, p < .4993); effect of interaction (F[4, 50] = 0.1004, p = .9818), n = 4 – 8 experimental replicates, n.s.).

To further characterize the functional pharmacology of these SV2-targeted compounds, we performed a series of *ex vivo* fast-scan cyclic voltammetry (FSCV) recordings in acute mouse striatal slices treated with PSL, UCB-F, or PRM **(Figure 3**). Extracellular dopamine (DA) transients evoked by a single-pulse (1p) electrical stimulation were monitored via fast-scan cyclic voltammetry (FSCV) as previously described^62^. To isolate the direct action of these ligands on dopamine release and reuptake, a comprehensive synaptic blocker cocktail was present continuously to eliminate indirect network-driven transmission mediated by glutamate, GABA, and acetylcholine receptors. We first evaluated the inhibitory profile of PSL on single pulse (1p)-evoked DA release. Bath application of PSL robustly suppressed DA release **(Figure 3A,B**), with a significant decreased of the peak amplitude of DA release apparent at a concentration of 10 µM (24.7 ± 5.8 %, P < 0.01), with maximum suppression at 100 µM (49.0 ± 5.5 %, P < 0.0001). Non-linear regression analysis of the concentration-response curve revealed that PSL inhibits evoked striatal DA release with a half-maximal inhibitory concentration (IC_50_) of 9.5 µM **(Figure 3C)**. Next, the effect of UCB-F was examined under identical experimental conditions. UCB-F likewise exhibited a dose-dependent inhibition of evoked DA transients **(Figure 3D,E)**, significantly reducing DA peaks at 50 (42.7 ± 4.3 %) and 100 µM (34.7 ± 2.2 %, P < 0.0001), with a calculated IC_50_ of 26.4 µM **(Figure 3F)**. In contrast to PSL and UCB-F, application of PRM at concentrations up to 200 µM had no effect on 1p-evoked DA release **(Figure 3G, H)**. We also observed that UCB-F, but not PSL or PRM, occasionally caused failure of evoked DA release after stimulation, suggesting that although there were no effects on FFN loading, UCB-F may affect other aspects of synaptic vesicle fusion and evoked neurotransmitter release.

**Figure 3.**
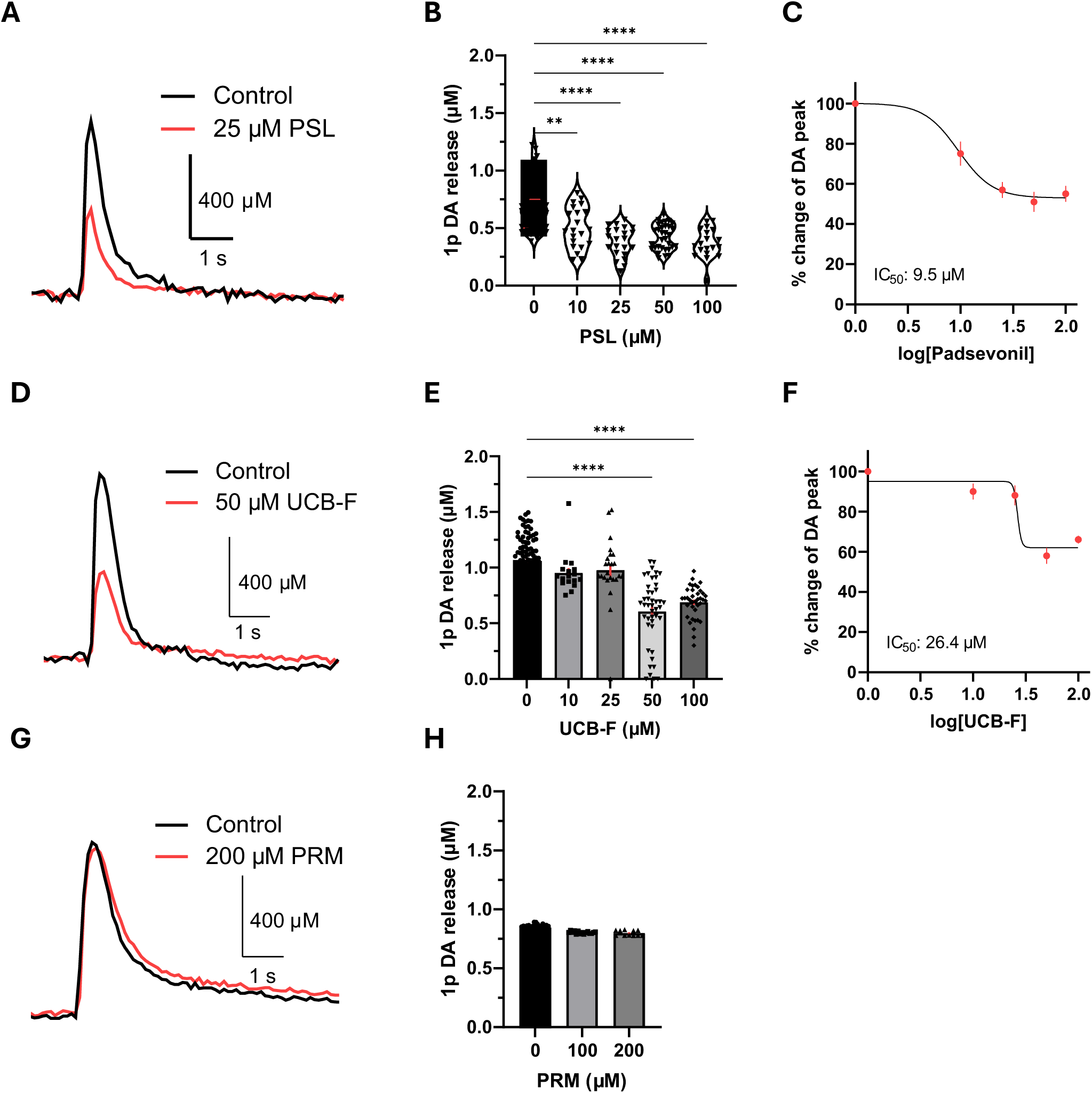
**A)** Representative traces of DA release in the dorsolateral striatum evoked by a single-pulse (1p) stimulation in the absence (Control, black) or presence of 25 µM padsevonil (PSL, red). **B)** Summary data showing dose-dependent inhibition of 1p-evoked DA release by PSL (Untreated: 56 sites/11 slices/5 mice; 10 µM: 21/4/3; 25 µM: 22/4/3; 50 µM: 32/6/4; 100 µM: 19/4/3). **C)** Dose-response curve of evoked DA release on PSL concentration (IC_50_ = 9.5 µM). Data are normalized to the control response and fitted with a non-linear regression curve. **D)** Representative traces of 1p-evoked DA release for control (black) and in the presence of 50 µM UCB-F (red). **E)** Dose-dependent effects of UCB-F on 1p-evoked DA release (Untreated: 88/15/6; 10 µM: 17/4/3; 25 µM: 24/4/3; 50 µM: 44/7/4; 100 µM: 38/5/3). **F)** Dose-response curve of UCB-F on the peak DA release (IC_50_ = 26.4 µM). **G)** Representative traces of 1p-evoked DA release for control (black) and in the presence of 200 µM primidone (PRM, red). **H)** Summary data showing the effect of PRM on evoked DA release (0 µM: 24/5/3; 100 µM: 15/4/3; 200 µM: 11/3/2). Data are presented as mean ± SEM. Each dots in the bar graph represent the recording site. **, **** p<0.01 or 0.0001 versus control by one-way ANOVA with Tukey’s post-hoc test.

### Cryo-EM of SV2C

Next, we sought to determine structures of SV2s bound to these ligands to better understand the molecular basis for their differences in affinity and selectivity. Previously, our group determined structures of SV2A and SV2B bound to the PSL derivative UCB2500, and PSL, respectively^42^. In those studies, we observed that thermostabilization induced by ligand binding made cryo-EM structure determination straightforward, and thus we first attempted to determine a structure of SV2C bound to PSL. We expressed full-length mVenus-tagged human SV2C in tsA201 cells, purified in the presence of PSL, and imaged by cryo-EM, resulting in a 3.6 Å reconstruction of the SV2C-PSL complex **(Figure 4A)**. Although this map provided sufficient detail to model protein-ligand interactions, reconstruction of the SV2C-PSL complex was challenging, owing to the high mobility of the luminal domain (LD), which hindered particle alignment. We attempted several strategies to add a fiducial to the LD, including botulinum neurotoxin A (BoNTA) and grafting of a previously identified SV2A-specific nanobody epitope onto the SV2C LD but could not overcome the high degree of LD flexibility^43^.

**Figure 4.**
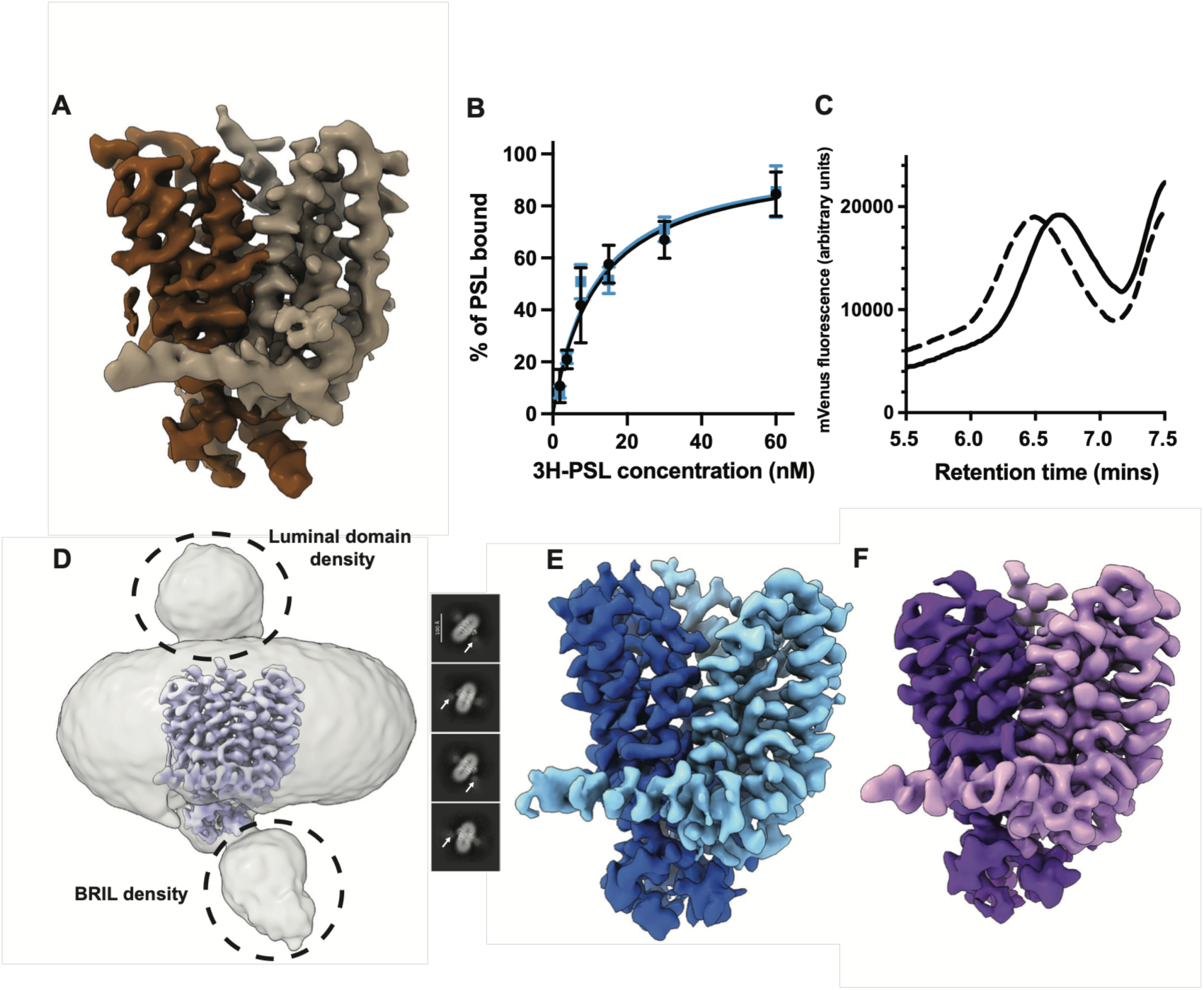
**A)** Cryo-EM reconstruction of SV2C bound to padsevonil, with N-terminal helix bundle density shown in dark brown and C-terminal helix bundle density shown in tan. **B)** Saturation binding of 3H-padsevonil to wild-type SV2C (black circles, K_d_ = 12 nM +/- 2 nM) and BRIL-SV2CΔ127 (blue squares, K_d_ = 11 nM +/- 2 nM). **C)** Fluorescence size exclusion chromatography trace of BRIL-SV2CΔ127 alone (solid) and BRIL-SV2CΔ127 with BAG2 Fab (dashed). **D)** Left: Gaussian-filtered cryo-EM map of UCB-F bound BRIL-SV2CΔ127 showing luminal domain and BRIL fiducial density; right: 2D classes of BRIL-SV2CΔ127 with arrows indicating diffuse BRIL fiducial density. **E)** Cryo-EM reconstruction of BRIL-SV2CΔ127 bound to UCB-F, with N-terminal helix bundle density shown in dark blue and C-terminal helix bundle shown in light blue. **F)** Cryo-EM reconstruction of apo BRIL-SV2CΔ127, with N-terminal helix bundle density shown in purple and C-terminal helix bundle density shown in pink.

Therefore, we added a fiducial on the intracellular domain (ICD) of SV2C instead. We truncated the first 127 residues of the disordered SV2C N-terminus, and added an N-terminal BRIL fusion protein linked directly to the ICD, allowing us to use a previously developed anti-BRIL Fab as a fiducial^63,64^. This construct, referred to as BRIL-SV2C-Δ127, retained binding to ^3^H-labeled PSL and addition of anti-BRIL Fab shifted the elution volume of the complex by fluorescence detection size exclusion chromatography (FSEC), indicating that the truncated SV2C remained functional and BRIL was accessible to the Fab **(Figure 4B and 4C)**. BRIL-SV2C-Δ127 was expressed and purified in large scale from tsA201 cells and used to prepare apo- and UCB-F-bound SV2C. The anti-BRIL Fab copurified with this construct after size-exclusion chromatography, and the Fab was clearly visible in raw cryo-EM micrographs **(Figure 4D, Extended Data 1B)**. Although Fab density was not resolved in our reconstructions, the additional BRIL feature attached to the ICD was sufficient to drive particle alignments, resulting in a 3.3 Å UCB-F bound SV2C reconstruction and a 3.5 Å apo SV2C reconstruction **(Figure 4E and 4F)**.

### Architecture of SV2C

Like SV2A and SV2B, SV2C adopts a canonical major facilitator superfamily fold, comprised of twelve transmembrane helices (TMs) arranged in two 6-TM inverted repeats forming a transmembrane domain (TMD) surrounding a central cavity. A large (approximately 100 amino acids) glycosylated loop connects TMs 7 and 8, forming the luminal domain (LD) of SV2C. Although we could not readily resolve the LD in our maps, crystal structures of this domain have previously been solved, showing that it adopts the pentapeptide repeat fold characteristic of the SV2A and SV2B LDs^29^. We could only partially resolve luminal loops 1 and 3 in our reconstructions, with interpretable density present for only the first non-helical residues. The ICD of SV2C contains several loops as well as a helix bundle formed by intracellular loops 1 and 3.

IL3 also contains an amphipathic helix that runs parallel to the membrane bilayer surface **(Extended Data 1C)**. In our reconstructions, the helical features of the ICD were readily resolvable. These features, conserved across all three SV2 family members, may be important for regulating synaptotagmin trafficking and transporter gating, and a human mutation in this region of SV2A has been linked to intractable epilepsy, suggesting an essential functional role^42,65^.

### Ligands bind a conserved primary site in SV2C

Consistent with other SV2 family members, an opening approximately 11.5 Å wide in the luminal side of SV2C leads to the primary ligand binding site located between the N- and C-terminal TM bundles **(Figure 5A-D)**. The 2-pyrrolidone core shared between all racetam SV2 ligands binds between Trp286 and Trp651 in SV2C. These residues are absolutely conserved amongst SV2s, analogous to Trp300/Trp666 in SV2A and Trp243/Trp607 in SV2B. Mutation of either Trp leads to a loss of ligand binding^42,44^. The SV2A double mutant W300A/W666A is sufficient to enable Syt trafficking in cultured neuronal cells, but not sufficient to restore aberrant electrical activity, suggesting that these residues play an important functional role that may be modulated by ligand binding^25^. Upon closer examination of the apo-SV2C cryo-EM map, we observed an unidentified density in the primary drug binding pocket, between Trp286 and Trp655 **(Extended Data 1D)**. A recently published apo-SV2A structure also reported unknown density in the primary drug binding site as well^45^. This suggests that an endogenous molecule present in cells or culture media binds SV2s and copurifies. Cryo-EM maps lack sufficient resolution and occupancy to identify these densities, and further work is needed to determine whether there are bona-fide endogenous SV2 ligands or transport substrates.

**Figure 5.**
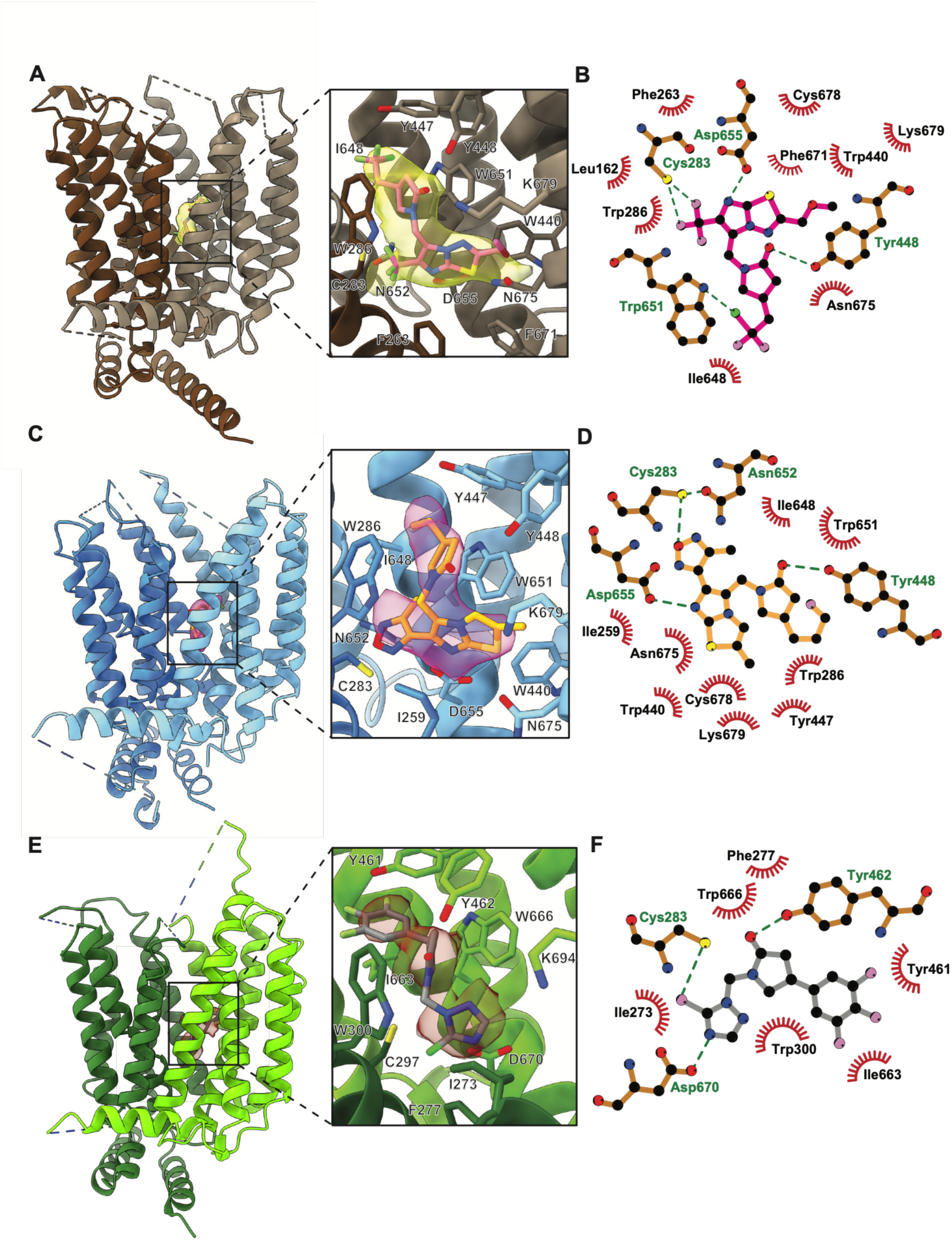
**A)** Model of SV2C bound to PSL, with cryo-EM density for PSL (pink sticks) highlighted in translucent yellow. **B)** Ligplot showing interactions between PSL (pink sticks) and SV2C (tan sticks), with hydrophobic contacts indicated by red lines and hydrogen bonds indicated by green lines. **C)** Model of SV2C bound to UCB-F, with cryo-EM density for UCB-F (orange sticks) highlighted in translucent yellow. **D)** Ligplot showing interactions between UCB-F (orange sticks) and SV2C (tan sticks), with hydrophobic contacts indicated by red lines and hydrogen bonds indicated by green lines. **E)** Model of SV2C bound to UCB-F, with cryo-EM density for PRM (grey sticks) highlighted in translucent brown. **F)** Ligplot showing interactions between PRM (grey sticks) and SV2A (tan sticks), with hydrophobic contacts indicated by red lines and hydrogen bonds indicated by green lines.

Positioned above the conserved tryptophan residues, a conserved dityrosine motif formed by Tyr447 and Tyr448 caps the ligands. Large aromatic residues commonly serve as gates in major facilitator superfamily transport proteins, and the dityrosine motif can be found in several related transport proteins with established transport functions, supporting the hypothesis that SV2s are transport proteins^66^. In this conformation, the dityrosine motif closes access to the central TMD region, making this structure a luminal-facing occluded conformation. A critical hydrogen bond between the hydroxyl group on the sidechain of Y448 and the 2-pyrrolidone carbonyl group is present in both the PSL and UCB-F structures **(Figure 5A-D)**. This interaction is important for binding of pyrrolidone derivatives, and ligands lacking the carbonyl at this position have poor affinity for SV2s^42^. PSL and UCB-F both share a large, fused ring system at the 1-position, a common feature of high-affinity SV2 ligands. This fused ring system points ‘downward’ into the cavity, oriented towards the intracellular side of SV2C. A conserved aspartate residue, D655, is oriented within hydrogen bonding distance of this ring system, and pKa calculations suggest that this residue is likely protonated in other SV2s, enabling it to serve as a hydrogen bond donor^42^. Mutation of the equivalent residue in SV2A and SV2B leads to a loss of binding affinity^42,44^. The remainder of the binding site is formed by weak, mostly hydrophobic interactions, and mutation of single residues in SV2A and SV2B does not have a strong effect on binding affinity, suggesting that each individual interaction only makes a small contribution towards overall ligand affinity^42,44^.

UCB-F shares the 2-pyrrolidone core and halogenated tail with other related high affinity ligands but also exhibits several key differences. The fused ring system at the 1-substituted position of UCB-F is an imidazothiazole ring, lacking a nitrogen atom present in the imidazothiadiazole ring of other racetam ligands. Instead of the trifluoromethyl substituent on the imidazothiazole ring, UCB-F contains a methylfurazan group at this position, and instead of the ether group located at the opposite end of the imidazothiazole ring, UCB-F has a methyl substituent. The 4-substitution from the central 2-pyrollidone core in UCB-F is a fluoropropyl group, while PSL possesses a difluorochloroethyl group, and UCB2500 has a trifluorobutyl group. Although UCB-F and PSL are comparable in size, the positions of large groups on each ligand are different, which causes UCB-F to occupy a slightly ‘higher’ position in the primary binding site.

The overall orientation of UCB-F is like PSL and UCB2500, with the fluoropropyl tail pointing upwards towards the lumen and the imidazothiazole ring system oriented towards the intracellular side of SV2C **(Figure 5C)**. The methylfurazan ring occupies the same space as the trifluoromethyl substituent of PSL, and is within 3.5 Å of the Cys283 sidechain, allowing for weak hydrogen bonding. Asn652 and Asn597 form a polar network with Cys283 which may further strengthen these interactions. To accommodate the interactions of this moiety with Cys283 and Asn652, the imidazothiazole ring shifts ‘upwards’ towards the luminal side by approximately 2 Å, with the rest of the ligand adjusting accordingly. Tyr448 maintains a hydrogen bond with the 2-pyrollidone core moiety, and the fluoropropyl substituent of UCB-F occupies approximately the same space as its equivalent groups in the UCB2500 and PSL-bound bound SV2A and SV2B structures^42,44^. The side chain of Asn644 is located over 5 Å from the fluoropropyl group of UCB-F, and no direct interactions can be made at this distance, thus this residue does not play an immediately obvious role in the selectivity of UCB-F.

### Plosaracetam binds the conserved primary site in SV2A

To further understand the structural basis of selectivity in the SV2 family, we determined the structure of SV2A bound to the experimental cognitive enhancer plosaracetam (PRM), which was tested in phase II clinical trials as a potential treatment for symptoms of Alzheimer’s disease, dementia, and other neurological conditions which cause cognitive decline^40,41^. We purified an N-terminally truncated SV2A construct, SV2AΔ64, from tsA201 cells in the presence of PRM, and imaged the complex with a previously discovered SV2A-specific nanobody, 8667-Nb, using cryo-EM^43^. This resulted in a 3.5 Å resolution reconstruction of the SV2A-PRM-8667-Nb complex **(Figure 5E, Extended Data 1E)**. Intriguingly, despite its reported lack of antiseizure effects, PRM binds the primary site similar to ASMs, and the overall protein conformation is nearly identical to previously reported LEV bound, BRV-bound, PSL bound and UCB2500-bound SV2A structures **(Figure 5E-F, Extended Data 1F)**^42,44,45^. In place of the halogenated tail on the 4-position of the pyrrolidone that is shared by other SV2 ligands, PRM has a bulky trifluorophenyl group, similar to the PET tracer molecule UCB-J. However, unlike UCB-J and other high-affinity ligands, PRM has a smaller chlorotriazole ring at the 1-position. The trifluorophenyl group occupies the same space as the halogenated tail of other ligands, orienting towards the luminal side of SV2A. The fluorine atoms of the trifluoromethyl group bind within 3 Å of the backbone carbonyl on Gly659, allowing for interactions between the electronegative fluorine atom and the partially positive backbone carbon, and the phenyl ring is also positioned to form π-stacking interactions with the sidechain of Tyr461. As observed in other pyrrolidone-bound SV2 structures, the sidechain hydroxyl group of Tyr462 forms a key hydrogen bond with the pyrrolidone-bound carbonyl oxygen. On the intracellular-facing side of the binding pocket, the chlorotriazole ring is positioned within hydrogen bonding distance of the conserved residue D670. The cryo-EM density for this moiety fits in two orientations, with the chlorine atom of the chlorotriazole ring facing either towards or away from the thiol group of Cys283, suggesting that PRM can bind in two poses **(Extended Data 1G)**. We hypothesized that the cysteine-facing pose would result in a higher-affinity interaction, and to test this possibility, we measured PRM binding to membranes expressing either wild-type or C297A-mutant SV2A. We found that mutation of this residue had no effect on PRM affinity, indicating that this interaction is dispensable for binding to SV2A **(Extended Data 1H)**.

### Determinants of SV2 ligand selectivity

Only two residues in the primary binding site of SV2s are not conserved. Cys283 in SV2C (equivalent to Cys297 in SV2A) is replaced by a glycine, Gly240, in SV2B, and Asn644 in SV2C is replaced by Gly659 in SV2A and Cys600 in SV2B. In all three PSL-bound SV2 structures, the sidechain of the conserved aspartate residue lies within 2.6 Å of the hydrogen bond acceptor atom on the imidazothiadiazole ring of PSL^42,44^. Alignment of ligand-bound SV2 structures reveals that PSL, UCB-F and PRM bind the primary site in similar poses, but subtle differences in positioning may be contributing to ligand selectivity **(Figure 6A)**. The sidechain of Cys297 in SV2A adopts a distinct rotamer, rotated 90° towards the cytosolic side of the protein relative to Cys283 in SV2C. Both rotameric configurations allow formation of a halogen bond with fluorine atoms in PSL. This interaction is not possible in SV2B due to a glycine at this position (Gly240), yet PSL still binds SV2B with high affinity, suggesting that this interaction is not a strong determinant of ligand specificity. Towards the luminal-facing end of PSL, the halogenated tail is oriented towards N644/G659 and lies within halogen bonding distance of the backbone carbonyl atom of these residues **(Figure 6B)**. Despite the presence of an additional potential halogen bond acceptor in SV2B, the halogenated tail of PSL does not come within close enough range of Cys600 to form any interactions, and mutation of Cys600 to glycine resulted in only a modest increase in PSL affinity, suggesting that this residue contributes minimally to the PSL affinity, which is independent of sidechain interactions^42^. Compared to SV2A and SV2B, the sidechains of the conserved tyrosine residues Tyr447 and Tyr448 in SV2C are shifted away from the primary binding site by approximately 1 Å and 1.3 Å, respectively. The distance between the sidechain hydroxyl group and the 2-pyrrolidone carbonyl is longest in SV2C at 3.6 Å, compared to 3.4 Å for SV2B and 3.1 Å for SV2A **(Figure 6B)**.

**Figure 6.**
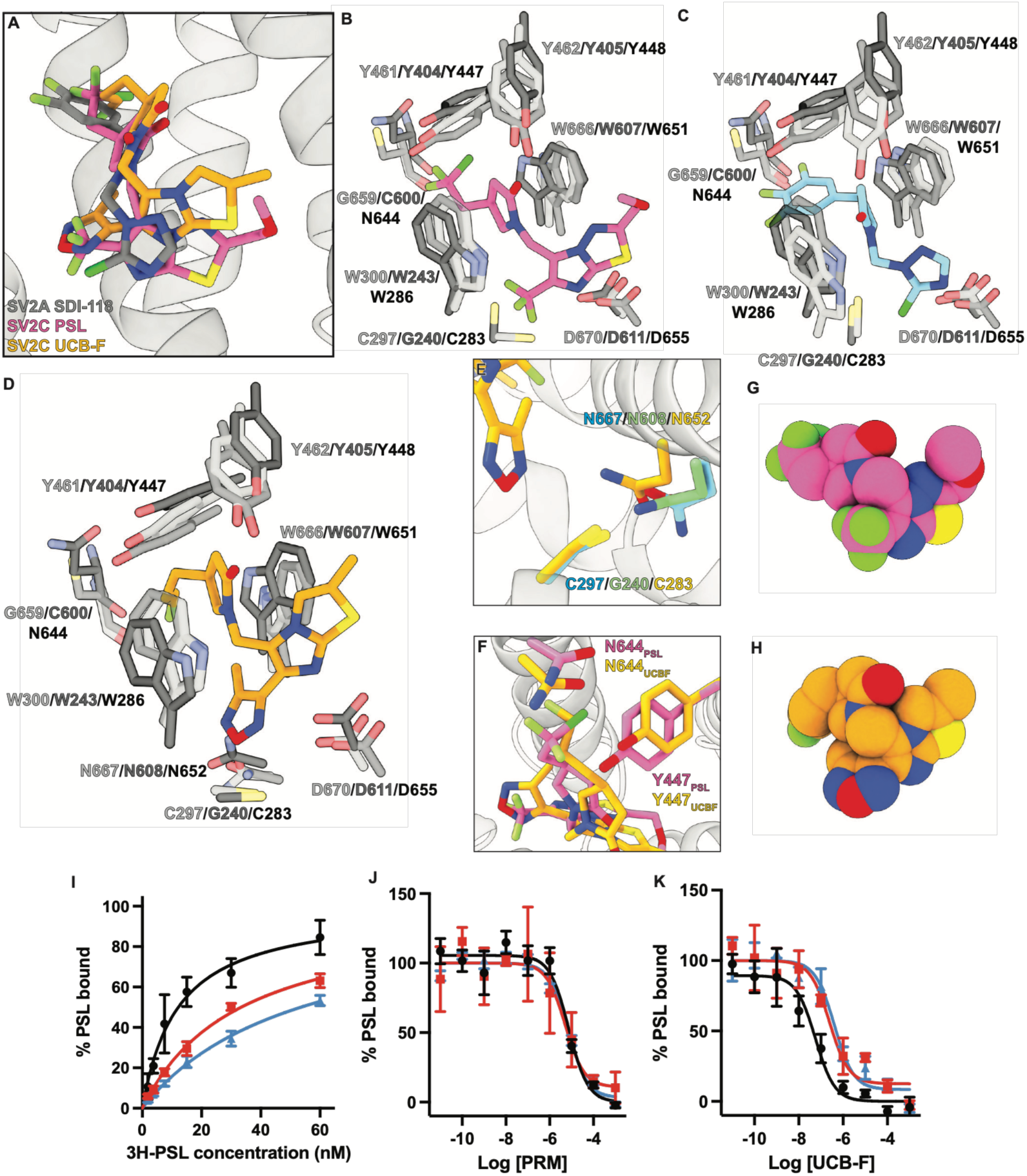
**A)** Alignment of SV2C-PSL, SV2C-, and SV2A-PRM structures. **B)** Alignment of SV2A-PSL (light grey sticks), SV2B-PSL (grey sticks), and SV2C-PSL (dark grey sticks, ligand shown as pink sticks) binding sites. **C)** Alignment of SV2A-PRM (light grey sticks, ligand shown as light blue sticks), SV2B-PSL (grey sticks), and SV2C-PSL (dark grey sticks, ligand shown as pink sticks) binding sites. **D)** Alignment of SV2A-PSL (light grey sticks), SV2B-PSL (grey sticks), and SV2C-UCB-F (dark grey sticks, ligand shown as orange sticks) binding sites. **E)** Comparison of interactions made by cysteine and asparagine sidechains in SV2A-PSL (blue sticks), SV2B-PSL (green sticks), and SV2C (orange sticks) around UCB-F (orange sticks). **F)** Comparison of Tyr447 and Asn644 movement between SV2C-PSL (pink sticks) and SV2C-UCB-F (orange sticks). **G)** Space-filling representation of PSL. **H)** Space-filling representation of UCB-F. **I)** Saturation binding of 3H-padsevonil to wildtype SV2C (black circles, K_d_ = 12 nM +/- 2 nM), Asn644Cys mutant SV2C (red squares, K_d_ = 33 nM +/- 2 nM), and Asn644Gly mutant SV2C (blue triangles, K_d_ = 53 nM +/- 6 nM). **J)** Competition binding of PRM with 5 nM 3H-padsevonil to wildtype SV2C (black circles, K_i_ = 4.9 uM +/- 1.1 uM), Asn644Cys mutant SV2C (red squares, K_i_ = 4.4 uM +/- 1.5 uM), and Asn644Gly mutant SV2C (blue triangles, K_i_ = 6.4 uM +/- 0.8 uM). **K)** Competition binding of UCB-F with 5 nM 3H-padsevonil to wildtype SV2C (black circles, K_i_ = 37 nM +/- 10 nM), Asn644Cys mutant SV2C (red squares, K_i_ = 212 nM +/- 77 nM), and Asn644Gly mutant SV2C (blue triangles, K_i_ = 368 nM +/- 91 nM).

Comparison of the SV2B-PSL and SV2C-PSL structures to the SV2A-PRM complex reveals that most binding site residues adopt a similar conformation, apart from Tyr462 in SV2A **(Figure 6C)**. The 2-pyrrolidone in the PRM core moves downward by approximately 1.6 Å compared to the equivalent group in PSL. To maintain the key hydrogen bond with the 2-pyrrolidone carbonyl, the side chain of Tyr462 moves downwards into the primary binding site by 1.6 Å relative to SV2B Tyr405 and 2.6 Å relative to SV2C Tyr448. We hypothesized that this movement could alleviate steric clashes between the rigid trifluorophenyl group in PRM and residues on TM10.

However, comparison with a previously determined SV2A structure in complex with UCB-J, which contains the same trifluorophenyl group in this position, showed that Tyr462 in these structures does not occupy the same position as the PRM bound state, thus the position of this residue is likely determined by another mechanism. Similar to PSL-bound structures, the trifluorophenyl group on PRM is also not close enough to interact with the non-conserved TM10 residue Gly659 in SV2A, equivalent to SV2B Cys600 and SV2C Asn644. However, SV2B lacks the additional cysteine in the binding site, equivalent to Cys297 in SV2A and Cys283 in SV2C, and this may be responsible for the 2-fold weaker K_i_ of PRM for SV2B vs. SV2C.

Next, we compared the UCB-F-bound SV2C structure to PSL-bound SV2A and SV2B structures **(Figure 6D)**. UCB-F is shifted approximately 1.7 Å towards the luminal side of SV2C relative to the position of PSL, and in order to accommodate this ligand position, Tyr447 and Tyr448 move ‘upwards’ by approximately 1 Å away from the primary binding site to maintain hydrogen bond distance to the 2-pyrrolidone carbonyl and the hydroxyl of Tyr448. This is accompanied by movement of Asp655 by 1.6 Å relative to its position in the PSL-bound structure, which hydrogen bonds with the imidazothiazole ring on UCB-F. The conserved residue Asn652 in SV2C, equivalent to Asn667 in SV2A and Asn608 in SV2B, appears to participate in a hydrogen bond network involving Cys283 and the methylfurazan ring on UCB-F. In SV2A, Asn667 points away from the primary site, and in SV2B Asn608 is located too far from these residues to participate in hydrogen bonding **(Figure 6E)**. The lack of these interactions in SV2A and SV2B likely partially contributes to the selectivity of UCB-F for SV2C. We also find that the sidechain of Asn644 shifts inward towards the center of the primary binding site by 1.4 Å in the UCB-F bound state relative to its position in the PSL bound state **(Figure 6F)**. This is consistent with the differences in the shape of each ligand, which have a comparable solvent accessible surface area but different arrangements of hydrogen bond donors and acceptors **(Figure 6G-H)**. Together, these rearrangements reduce the distance between the sidechain carbonyl of Asn644 and the sidechain hydroxyl group of Tyr447 from 4.6 Å in the PSL-bound structure to 3.2 Å in the UCB-F structure, allowing for the formation of a new hydrogen bond in this configuration. SV2A, which possesses a glycine at this position, cannot form this interaction, and the equivalent cysteine residue in SV2B, Cys600, is also 4.6 Å away from the equivalent Tyr404 sidechain, which prevents hydrogen bond formation. Recently, molecular dynamics work using a homology model of SV2C predicted that Tyr447 and Asn644 could form a persistent hydrogen bond, and that this hydrogen bond could stabilize UCB-F binding to the primary site^67^. Thus, we hypothesized that these residues could be involved in controlling access and selectivity at the primary site of SV2C. We mutated Asn644 in SV2C to the corresponding residues found in SV2A and SV2B at this position and investigated binding of PSL, PRM, and UCB-F. Mutation of Asn644 decreased PSL affinity from 12 nM to 53 nM for Asn644Gly and to 33 nM for Asn644Cys **(Figure 6I).** We then performed competition binding experiments with UCB-F and PRM using ^3^H-labeled PSL. In both cases, mutation of Asn644 had little impact on PRM affinity, with a Ki of 6.4 µM for Asn644Gly and 4.5 µM for Asn644Cys, compared with a Ki of 4.9 µM for wild-type SV2C **(Figure 6J)**. The K_i_ of UCB-F decreased from 37 nM in wild-type SV2C to 368 nM for Asn644Gly and 212 nM for Asn644Cys **(Figure 6K)**.

Although these mutations weakened UCB-F binding by approximately 10-fold, we could not fully recapitulate the 20-fold weaker binding of UCB-F to SV2A, or its lack of binding to SV2B. The differences in binding affinity for these SV2C mutants vs. SV2A and SV2B can be explained by two factors: because of the orientation of Asn667 in SV2A, it cannot form a hydrogen bond network with Cys297 as seen in the UCB-F-bound SV2C model; and SV2B lacks a sidechain at Gly240 (analogous to Cys283 in SV2C) which cannot form any interactions with UCB-F. Curiously, the affinity of PRM for SV2C did not increase by mutating Asn644 to glycine, despite this being the only non-conserved residue comparing the SV2A and SV2C primary binding sites. This suggests that although Asn644 plays a role in the selectivity of UCB-F for SV2C, a different mechanism underpins the selectivity of PRM for SV2A.

Comparing apo-SV2C and UCB-F bound SV2C to the SV2C-PSL complex structure, we observed that the upper luminal-facing half of TM7 has a different conformation. The cytoplasmic-facing half of TM7 in these structures adopts a similar conformation around the cytoplasmic side of the primary binding site. Beginning at the conserved residue Phe445, close to the conserved gate residues Tyr447 and Tyr448, TM7 in the apo- and UCB-F bound structures begins to pitch away from the primary site by approximately 8° relative to the same region in the PSL-bound state **(Figure 7A)**. We compared this region of SV2C to other SV2 structures **(Extended Data 3A-B)**. When SV2A is bound to ligands, TM7 adopts a closed conformation to varying degrees, with the most closure observed in models where SV2A is bound to both a primary site ligand and the allosteric ligand, UCB1244283. PRM, UCB-J, UCB2500, and LEV also induce closure of TM7. The conformation of TM7 in the PSL-bound state resembles an intermediate between the apo-SV2A structure, which has the most open conformation, and the closed UCB-J structure. Interestingly, SV2B TM7 in both the apo- and PSL-bound states is closed. Movements of the transmembrane region that lines the primary binding site can provide allosteric control over binding of molecules at the primary site, and similar mechanisms are likely important for binding of endogenous molecules to SV2s.

**Figure 7.**
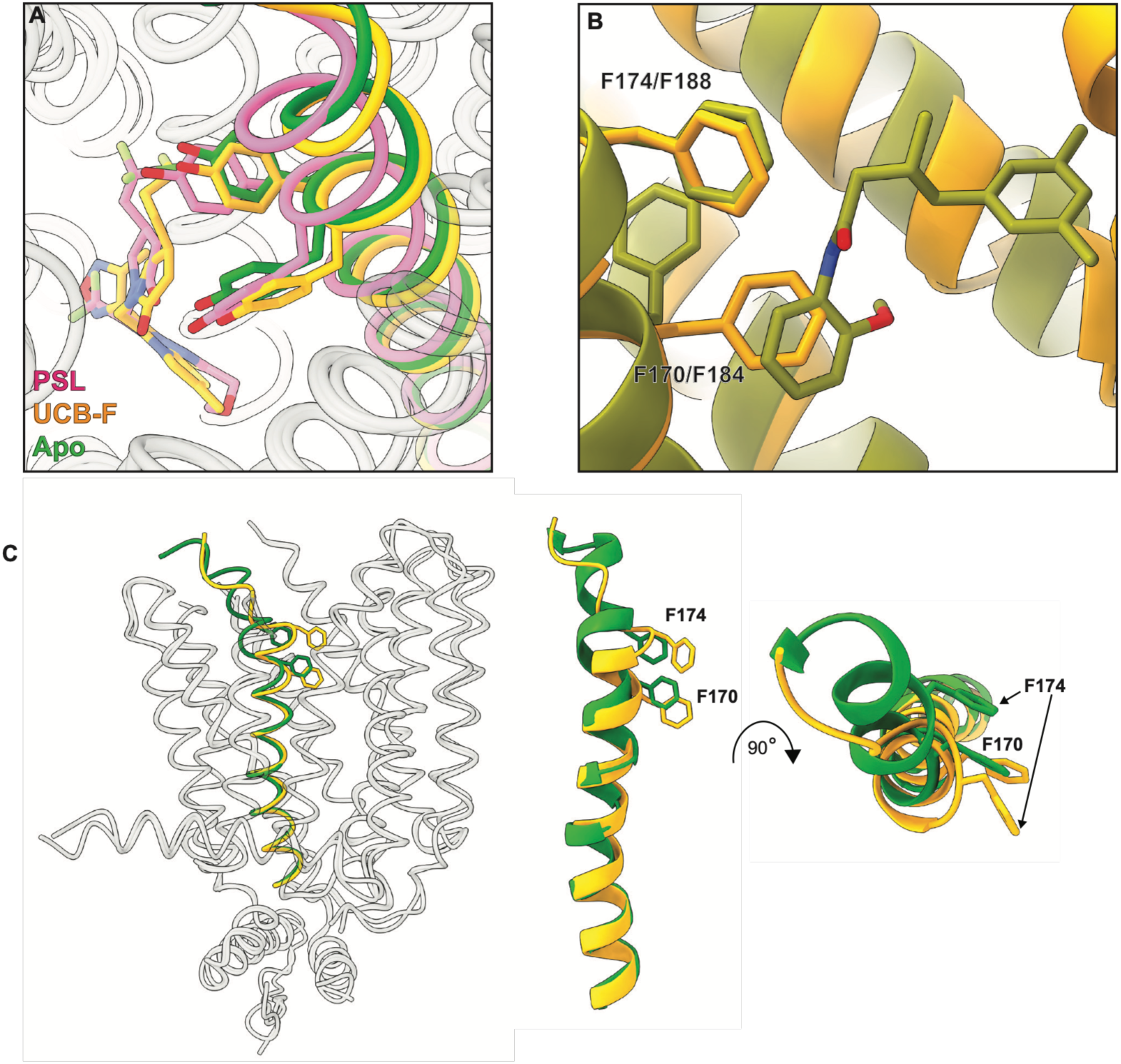
**A)** Movement of TM7 in SV2C between the apo- (green), UCB-F- (orange), and PSL-bound (pink) states. **B)** Comparison of the allosteric modulator binding site between SV2A-BRV-UCB1244283 (PDB 9NTC, olive green with ligand shown as olive green sticks) and UCB-F-bound SV2C (orange). **C)** Movement of TM1 between apo (green) and UCB-F-bound SV2C (orange).

We also found subtle differences in the position of TM1 in SV2C compared to other SV2s. In our PSL-bound SV2C reconstruction, we observed that the luminal half of TM1 had interpretable main chain density, but weak density for the large aromatic sidechains of Phe170 and Phe174 **(Extended Data 3B-C)**. These phenylalanine residues are conserved amongst SV2s and have been implicated in allosteric modulation of primary site ligand binding in SV2A^43,44^. The absence of sidechain density for these residues suggests that in the PSL-bound state, this region of SV2C is dynamic. In the apo and UCB-F bound reconstructions, we could readily model density for the sidechains of these residues **(Figure 7B, Extended Data 3D-E)**. In the UCB-F model, TM1 most closely resembles the “closed” conformation observed in SV2A bound to high-affinity primary site ligands, as well as the ternary complex formed with BRV and the allosteric modulator compound UCB1244283^42,43^. This is consistent with the hypothesis that closure of the conserved phenylalanines over the primary binding site drives high affinity ligand binding. However, the position of Phe170 in SV2C is different than the analogous residue, Phe184, in SV2A. In SV2A, this residue points ‘upwards’ towards the luminal side of the protein, away from the allosteric modulator binding site. In SV2C, this residue orients ‘downward’ toward the primary ligand binding site, approximately 3 Å away from the analogous residue in SV2A, causing it to overlap with the allosteric modulator binding site in SV2A. This steric hindrance combined with other residue substitutions in the allosteric site between SVA and SV2C, such as the replacement of Gly659 with Asn644, likely could prevent binding of UCB1244283. In the apo SV2C reconstruction, we find that TM1 occupies a different conformation in its luminal half than our ligand-bound structures. Phe170 occupies a similar position to the UCB-F bound SV2C structure, but beginning at Val171, the remainder of TM1 bends outwards away from the primary site. This causes the position of Phe174 to rotate approximately 45° counterclockwise from the “closed” position that is observed in other SV2 structures **(Figure 7C)**. Large aromatic amino acids frequently act as gates in transport proteins, and these dynamic residues likely play a key role in the function of SV2C.

## Discussion

The precise biological function of SV2s, and the mechanism by which pyrrolidone-class ASMs exert their effects still remains to be fully elucidated despite decades of research. The effects of BRV and LEV on neurotransmission have been studied in-vitro and in-vivo, but these drugs are selective for SV2A, and thus little is known about the effects of modulating SV2C function on the brain^68–72,72–74^. Our FFN206 uptake studies in cells show that there are only minor effects on loading when cells expressing VMAT2 and SV2C are treated with PSL and no effect is observed with UCB-F or PRM. In contrast, our ex-vivo studies show that treatment of brain slices with PSL or UCB-F causes a decrease in evoked dopamine release. However, the concentrations required to observe an inhibitory effect on dopamine release from brain slices were several orders of magnitude higher than the binding affinity of these compounds for SV2C. We note that for brain slice experiments, compounds were incubated with 250 µm thick brain slices for 2-3 hours at room temperature, which may limit accessibility. Thus, local concentrations of these compounds in the brain slice may have been significantly lower than their concentrations in the surrounding solution.

Second, our FSCV data taken together with our FFN206 uptake data suggests that these compounds impact vesicular release with little impact on vesicular loading, and because of this, we cannot rule out a mechanism involving both SV2C and Syt. All SV2s are known to bind to and traffic Syts, which play a critical role in modulating fusion of synaptic vesicles. It has been speculated that ASMs exert their effects by affecting SV2-Syt complex formation or function. Previously, we found that complex formation was not affected by the presence of ligands, and that the affinity of PSL for the SV2A-Syt1 complex was unchanged^42^. Because our assays only measure binding affinity and complex formation, we cannot rule out effects on function of Syt or the SV2-Syt complex that are mediated by these compounds. Furthermore, several other factors such as membrane composition and calcium concentration are also known to regulate Syt function. Thus, the IC50 of these compounds on dopamine release in our ex-vivo experiments may differ from the in-vitro SV2C binding affinity. However, we cannot rule out compound binding to other targets as a confounding factor in these results. PSL was designed to bind SV2s with high affinity while maintaining lower affinity for the benzodiazepine site on the GABA_A_ receptor, where it acts as a partial agonist with an affinity in the micromolar range^34^. Although a blocking cocktail containing 25 µM of the GABA_A_ antagonist picrotoxin was used throughout these experiments, the decrease in evoked release may also be caused by potentiation of presynaptic GABA_A_ receptors by PSL, leading to suppression of release, a well-known function of presynaptic GABA receptors. Furthermore, the high concentrations of PSL and UCB-F that were required to induce this effect may also be a result of engagement of other unknown low-affinity binding sites in neurons that impact release. Thus, while these results are interesting and may suggest that they are mediated through SV2C-Syt complexes, we interpret them with caution.

Until recently, understanding the basis of SV2 ligand selectivity has been hindered by a lack of structural information about the protein family. The non-conserved cysteine residue Cys297 in SV2A has previously been implicated in the selectivity of LEV, but little is known about the specificity of high-affinity ligands^42^. Our structures show that despite the high degree of sequence conservation, small but functionally important differences in the arrangement of residues in the primary binding site underpin the selectivity of racetam ligands. PSL, which binds all three SV2 family members with high affinity, binds to SV2C in the same pose as other SV2s, and forms similar interactions between conserved tyrosine and tryptophan residues that have previously been shown to be important for racetam binding. UCB-F, which binds SV2A approximately 20-fold weaker than SV2C, and does not appreciably bind SV2B, also interacts with these residues. The addition of a methylfurazan ring engages Cys283, Asn652 and Asn597 are also likely important for these interactions as they are within hydrogen bonding distance of Cys283 but don’t directly engage with UCB-F. These residues are conserved in SV2A, but the equivalent asparagine residue to Asn652 in SV2A is Asn667 which faces away from the ligand in other SV2A structures and cannot participate in hydrogen bonding, which likely weakens the interactions with this moiety. SV2B does not have a sidechain at the equivalent position to Cys283, Gly240, to form this hydrogen bond network. Interactions with Cys283 and Asn652 also cause UCB-F to occupy a slightly different position in the primary site compared to PSL, where Asn644 is within hydrogen bonding range of Tyr447. This residue is part of the conserved dityrosine motif found in all SV2s, and these residues are involved in both direct interactions with primary site ligands through hydrogen bonding with the 2-pyrrolidone carbonyl group, as well as occlusion of the primary site. Additional hydrogen bonding to Tyr447 likely has a stabilizing effect on the outward-facing occluded conformation observed in our structures, and drives the high affinity binding of UCB-F to SV2C. SV2A has a glycine at this position which cannot form any hydrogen bonds, and SV2B has a cysteine at this position with a smaller sidechain that cannot reach Tyr404 to form the analogous hydrogen bond. Indeed, mutation of Asn644 to either glycine or cysteine decreased the K_i_ of UCB-F for SV2C by 10-fold, supporting the hypothesis that stabilization of the occluded conformation is important for selective high-affinity binding to SV2C. The lack of interactions between Asn667 and Cys297 in SV2A can explain why Asn644Gly mutant SV2C still does not bind UCB-F with the same affinity as SV2A, and Gly240 in SV2B at this position explains why Asn644Cys mutant SV2C is able to bind UCB-F with relatively high affinity, compared to negligible binding observed in our competition experiment with SV2B. The selectivity of PRM appears to be mediated through a different mechanism. Mutation of Asn644 to glycine or cysteine did not have significant effects on the K_i_ of PRM for SV2C, despite the Asn644Gly SV2C mutant primary site having an identical sequence to SV2A. Furthermore, mutation of Cys297 to alanine in SV2A did not have appreciable effects on PRM binding affinity either, suggesting that the substitution of cysteine with glycine at this position in SV2B is also not responsible for the weaker binding of PRM.

Several structures of SV2A and SV2B have been published recently in complex with a variety of ligands, allowing us to perform detailed comparisons of the positions of TM1 and TM7 in the primary binding site^30,42–44^. TM7 has been implicated in allosteric control of the SV2A primary binding site, through interactions with both the allosteric modulator compound UCB1244283 and high-affinity ligands, such as PSL and UCB-J^43,44^. Movement of conserved phenylalanine residues on the luminal side of the protein, located above the dityrosine motif, further occludes the primary site and increases the affinity of bound ligands. We found that in the PSL-bound state, the backbone of SV2C TM1 could be modelled in our cryo-EM density, but the sidechains for Phe170 and Phe174 could not be reasonably modelled, suggesting that there was significant disorder in this region. In the UCB-F bound state, Phe170 and Phe174 were readily resolvable, and resembled the “closed” conformation that was previously observed in structures of SV2A bound to high-affinity ligands. Importantly, the sidechain of Phe170 is in a different position than the analogous residue in SV2A, Phe184, and comparison of the two structures shows that Phe170 would directly overlap with the UCB1244283 binding site. Combined with other steric hindrance caused by substitution of Gly659 in SV2A, which makes up one face of the allosteric modulator binding site, with Asn644 in SV2C, this suggests that UCB1244283 does not bind to SV2C. In our apo reconstructions of SV2C, TM1 faced outwards from the transmembrane domain, widening the luminal opening. We also note that TM7, which connects to the N-terminus of the luminal domain in SV2s, also showed subtle differences in positioning that appear to be induced by ligands. In SV2A, TM7 was most “closed” in the BRV-UCB1244283-bound state, with less closure observed in the LEV, PRM, UCB2500, and UCB-J complexes. Apo-SV2A and PSL-bound SV2A had the most “open” TM7 conformation, while apo- and PSL-bound SV2B both occupied a “closed” conformation. Apo- and UCB-F bound SV2C exhibited the most “open” TM7 conformation out of all the compared SV2 structures, and SV2C-PSL TM7 closely aligned with TM7 in SV2A-PSL and SV2B-PSL. The conformation of TM7 does not show any trend with increasing ligand affinity, but these small conformational differences may be another important factor for determining selectivity of SV2 ligands and are likely also involved in control of endogenous small molecule binding to SV2s.

Taken together, our work shows that PSL and the SV2C-selective ligand UCB-F effect dopamine release through a mechanism that is distinct from the inhibition of packaging into synaptic vesicles. UCB-F can selectively bind to SV2C by engaging additional residues around the binding site to increase the stability of the outward-facing occluded conformation that is observed in racetam-bound SV2 structures. This work provides a template for rational drug design for ligands specifically targeting SV2C, and may be useful for developing future treatments for epilepsy as well as dopaminergic neurological conditions such as PD.

## Extended data figure legends

**Extended Data Figure 1.** **A)** Competition binding of UCB-J with 20 nM 3H-padsevonil to SV2C-enriched membranes. K_i_ = 750 nM +/- 107 nM. **B)** Size exclusion chromatography of BRIL-SV2CΔ127 bound to BAG2. **C)** SDS-PAGE gel of BRIL- SV2CΔ127 samples. Before 3C addition, BRIL-SV2CΔ127 is fused to an mVenus tag and runs at a higher molecular weight on the gel. SV2C bands (*) are diffuse due to glycosylation of the luminal domain. BAG2 appears as a band at 20 kDa (#). C) Architecture of SV2C showing intracellular helical bundle, conserved amongst SV2s. The N-terminal helix ILH1 (blue) forms a bundle with ILH3a and ILH3b (light blue). ILH3c (purple) runs parallel to the membrane bilayer surface. **D)** An unidentified density (dark blue) is found between W286 and W651 (grey sticks) in the apo-SV2C cryo-EM reconstruction. Density corresponding to amino acid sidechains is shown as a transparent blue surface. **E)** Cryo-EM reconstruction of SV2A bound to PRM, with N-terminal helices shown in dark green and C-termina helices shown in light green. **F)** Overlay of SV2A-PRM model with SV2A-PSL (pink, PDB 9OKJ), SV2A-UCB2500 (brown, PDB 8UO9), LEV (light blue, PDB 9OKG) and BRV (yellow, PDB 8K77). The conformation of PRM-bound SV2A is similar to the conformation of SV2A bound to other ligands. **G)** The cryo-EM density for PRM (transparent brown surface) allows the chlorotriazole ring to be fit in two orientations, facing towards Cys297 (pink sticks, Cys297 as green sticks) or away from Cys297 (light blue sticks). **H)** Binding of 3H-PRM to wild-type SV2A (black circles, K_d_ 14 nM +/- 1 nM) or C297A mutant SV2A (red squares, Kd 12 nM +/- 1 nM).

**Extended Data Figure 2.** Treatment of HEK-VMAT-SV2C cells with PSL (black bars), PRM (pink bars), or UCB-F (teal bars) causes a dose-dependent increase in expression of SV2C relative to DMSO.

**Extended Data Figure 3.** **A)** Comparison of TM7 in apo-SV2A (brown, PDB 9OKF) to BRV-modulator-SV2A (gold, PDB 9NTC), LEV-modulator-SV2A (blue, PDB 9PRS), PRM-SV2A (red), UCB-J-SV2A (purple, PDB 9PCB), UCB2500-SV2A (pink, PDB 8UO9), PSL-SV2A (light blue, PDB 9OKJ), and LEV-SV2A (black, PDB 9OKG). **B)** Comparison of TM7 in apo-SV2B (green, PDB 9F1R) to PSL-SV2B (pink, PDB 8UO8). **C)** Poor density for aromatic sidechains of Phe170 and Phe174 on the luminal half of TM1 in the cryo-EM reconstruction of SV2C-PSL (transparent blue surface). The backbone could be fit for these residues (grey sticks) but no sidechain density was present. **D)** Comparison of TM1 Phe170 and Phe174 cryo-EM density in the apo-SV2C reconstruction (transparent blue surface, model shown as grey sticks). **E)** Comparison of TM1 Phe170 and Phe174 cryo-EM density in the SV2C-UCB-F reconstruction (transparent blue surface, model shown as grey sticks).

**Extended Data Figure 4.** Cryo-EM data collection and processing of SV2C-PSL. A representative micrograph and 2D class averages are shown. 46,200 micrographs were collected, and template picking yielded approximately 10.5 million particles. After an initial volume was generated with 2D classification and ab-initio reconstruction, several rounds of heterogenous refinement in Cryosparc against a starting SV2C volume and several decoy volumes consisting of empty micelles and debris were used to clean up the particle stack. Particles with good SV2C features were reextracted at full box size and subjected to an HR-HAIR ab-initio reconstruction to generate a higher quality SV2C volume. This volume was used for non-uniform refinement, yielding a 5 Å reconstruction with approximately 137,000 particles. This volume was used for extensive 3D classification and Blush refinement in Relion, followed by CTF refinement and Bayesian polishing. Local refinements of the transmembrane domain produced a final reconstruction of 70,940 particles at 3.56 Å resolution.

**Extended Data Figure 5.** Interpretation of the SV2C-PSL cryo-EM reconstruction. Fits for all 12 transmembrane helices and the intracellular domain are shown, as well as local resolution of the SV2C-PSL map, Euler angle distribution of the reconstruction, FSC curve of the SV2C-PSL map, and the map-to-model FSC correlation of the SV2C-PSL model to the SV2C-PSL map.

**Extended Data Figure 6.** Cryo-EM data collection and processing of apo-BRIL-SV2CΔ127. A representative micrograph and 2D class averages are shown. 36,766 micrographs were collected, and template picking yielded approximately 4.4 million particles. After an initial volume was generated with 2D classification and ab-initio reconstruction, several rounds of heterogenous refinement in Cryosparc against a starting SV2C volume and several decoy volumes consisting of empty micelles and debris were used to clean up the particle stack. Particles with good SV2C features were reextracted at full box size and subjected to an HR-HAIR ab-initio reconstruction to generate a higher quality SV2C volume. This volume was used for non-uniform refinement, yielding a 3.86 Å reconstruction with approximately 294,000 particles. This volume was used for 3D classification and CTF refinement in Relion. Local refinements of the transmembrane domain produced a final reconstruction of 47,041 particles at 3.5 Å resolution.

**Extended Data Figure 7.** Interpretation of the apo-BRIL-SV2CΔ127 cryo-EM reconstruction. Fits for all 12 transmembrane helices and the intracellular domain are shown, as well as local resolution of the apo-BRIL-SV2CΔ127 map, Euler angle distribution of the reconstruction, FSC curve of the apo-BRIL-SV2CΔ127 map, and the map-to-model FSC correlation of the apo-BRIL-SV2CΔ127 model to the apo-BRIL-SV2CΔ127 map.

**Extended Data Figure 8.** Cryo-EM data collection and processing of BRIL-SV2CΔ127-UCB-F. A representative micrograph and 2D class averages are shown. 20,982 micrographs were collected, and template picking yielded approximately 4 million particles. After an initial volume was generated with 2D classification and ab-initio reconstruction, one round of heterogenous refinement in Cryosparc against a starting SV2C volume and several decoy volumes consisting of empty micelles and debris was used to clean up the particle stack. Particles with good SV2C features were reextracted at full box size and subjected to an HR-HAIR ab-initio reconstruction to generate a higher quality SV2C volume, followed by several more rounds of heterogenous refinement to further remove bad particles. This particle stack was used for non-uniform refinement against an ab-initio volume, yielding a 4.5 Å reconstruction with approximately 72,000 particles. This volume was used for 3D classification and Blush refinement in Relion, followed by CTF refinement and Bayesian polishing. Local refinements of the transmembrane domain produced a final reconstruction of 40,132 particles at 3.29 Å resolution.

**Extended Data Figure 9.** Interpretation of the BRIL-SV2CΔ127-UCB-F cryo-EM reconstruction. Fits for all 12 transmembrane helices and the intracellular domain are shown, as well as local resolution of the BRIL-SV2CΔ127-UCB-F map, Euler angle distribution of the reconstruction, FSC curve of the BRIL-SV2CΔ127-UCB-F map, and the map-to-model FSC correlation of the BRIL-SV2CΔ127-UCB-F model to the BRIL-SV2CΔ127-UCB-F map.

**Extended Data Figure 10.** Cryo-EM data collection and processing of SV2A-8667Nb-PRM. A representative micrograph and 2D class averages are shown. 20,982 micrographs were collected, and template picking yielded approximately 4 million particles. After an initial volume was generated with 2D classification and ab-initio reconstruction, several rounds of heterogenous refinement in Cryosparc against a starting SV2A volume and several decoy volumes consisting of empty micelles and debris was used to clean up the particle stack. Particles with good SV2A features were reextracted at full box size and subjected to an HR-HAIR ab-initio reconstruction to generate a higher quality SV2A volume. This volume was subjected to non-uniform refinement, yielding a 4.3 Å reconstruction with approximately 85,000 particles. This volume was used for 3D classification and Blush refinement in Relion, followed by CTF refinement and Bayesian polishing. Local refinements of the transmembrane domain produced a final reconstruction of 26,282 particles at 3.53 Å resolution.

**Extended Data Figure 11.** Interpretation of the SV2A-8667Nb-PRM cryo-EM reconstruction. Fits for all 12 transmembrane helices and the intracellular domain are shown, as well as local resolution of the SV2A-8667Nb-PRM map, Euler angle distribution of the reconstruction, FSC curve of the SV2A-8667Nb-PRM map, and the map-to-model FSC correlation of the SV2A-8667Nb-PRM model to the SV2A-8667Nb-PRM map.

## Methods

### Cell culture

HEK293 cells were stably transfected with pcDNA3.1 vectors (Life Technologies) containing human vesicular monoamine transporter 2 (VMAT2; HEK-VMAT2) utilizing Zeocin selection, and human synaptic vesicle glycoprotein 2C (SV2C; HEK-VMAT2-SV2C) utilizing geneticin selection. Culture media consisted of Dulbecco’s Modified Eagle Medium (DMEM) with 4.5 g/L glucose (Corning), 10% fetal bovine serum (Fisher), .5% penicillin–streptomycin (Sigma), and 100 mg/mL Zeocin (Fisher), with the addition of 250 ug/mL Geneticin (Fisher) for double-stable HEK-VMAT2-SV2C culturing. All HEK293 cells were maintained in an incubator at 37C with 5% CO_2_ on 10-cm cell culture dishes coated with poly-D-Lysine (Sigma). tSA-201 cells were cultured in suspension in SFM4-Transfx-293 media (Cytiva) containing 1% FBS (Sigma) and 4 mM L-glutamine (Sigma). For adherent transfections, tSA-201 cells were grown in DMEM containing 10% FBS. All tSA-201 cells were maintained at 37C with 5% CO_2_. Sf9 cells were cultured in SIM SF media (SinoBio) at 27C in shaker flasks.

### Fluorescent False Neurotransmitter Uptake Assays

FFN206 uptake assays were performed in a modified fashion as previously described.^60,75^ Briefly, HEK-VMAT2 and HEK-VMAT2-SV2C were seeded at 40,000 cells per well on a PDL-coated black-walled, clear-bottomed, half-area, 96-well plate (Corning). 24 hours after seeding, culture media was removed, and cells were treated with DMSO (0.05% or 0.1%), Padsevonil (MedChemExpress), PRM (Pharmaron), or UCB-F (Pharmaron), at 1uM, 10uM, 25uM, or 50uM in DMEM (4.5 g/L glucose, sodium pyruvate; without L-glutamine, phenol red). Following 2h of incubation with compounds, FFN206 was added to a final concentration of 2uM for 1 hour. Following FFN206 incubation, drugs and FFN206 were removed and replaced with a 1:25 trypan blue solution in DMEM (4.5 g/L glucose, sodium pyruvate; without L-glutamine, phenol red). Plates were analyzed using a BioTek Synergy H1 multiplate reader at 37C using end-point fluorescent intensity measurements with excitation λ = 369 and λ = 464. Fluorescent values were corrected by subtracting background fluorescent values obtained from cells on the same plate with the same treatments without FFN206. For uptake experiments, each 96-well plate contained 9 technical replicates per cell line:treatment group. An average for the control condition (HEK-VMAT2:DMSO; HEK-VMAT2-SV2C:DMSO) was calculated, and each well value was expressed as a percent of this control. A single averaged value was calculated for each cell line:treatment group to represent an experimental replicate. Experimental replicates were combined across three to four separate experiments in which one HEK-VMAT2 plate and two HEK-VMAT2-SV2C plates were evaluated. Plates were observed at 4x on an APX100 Upright Fluorescent Microscope (Evident) to confirm cell adherence and no evidence of aberrant fluorescence. Wells were excluded due to pipette error, aberrant fluorescence due to contamination, or cell detachment if necessary. Two-Way ANOVA with Šídák’s multiple comparisons test was performed using GraphPad Prism10 to determine if the dose of compound had a significant impact on uptake within the individual cell line, and to determine if each dose of compound had a significant impact on uptake compared to the other cell line.

### Protein expression experiments

HEK-VMAT2-SV2C cells were seeded at 750,000 cells per well on a PDL-coated 6-well plate (Corning). 24 hours after seeding, culture media was removed and cells were treated with DMSO (0.1% or 0.2%), PSL, PRM, or UCB-F at 0.1uM, 1uM, 10uM, 50uM, or 100uM in DMEM (4.5 g/L glucose, sodium pyruvate; without L-glutamine, phenol red). 48 hours after treatment, cells were collected for Western blot analysis. Cells were rinsed with cold PBS and lysed in M-PER lysis buffer (Thermo) for 10m on ice. Lysates were briefly sonicated and centrifuged at 14,000 rpm for 15m at 4°C. Supernatant was collected, separated on 4–12% Bis-Tris gels (Invitrogen), and transferred onto nitrocellulose membranes using a Mini Blot Module (Invitrogen) at 17V for 70m. Membranes were blocked with 5% nonfat dry milk. Primary antibodies SV2C (Proteintech, 21271-1-AP; 1:1,500) and actin (Cell Signaling Technology, 12748S; 1:3,000); secondary antibody IRDye 800 (Licor, 926-32211; 1:15,000). Membranes were scanned using a Licor Odyssey CLx with quantification performed in Licor Image Studio.

### Fast scan cyclic voltammetry (FSCV)

FSCV recordings were performed in acute coronal striatal slices (250 μm thick) obtained from 3 month-old C57BL/6 mice. After brain slicing, the tissue was allowed to recover in normal artificial cerebrospinal fluid (ACSF) continuously bubbled with 95% O2 / 5% CO2 for 1 hour at 34 °C. Slices were then transferred to isolated incubation chambers and incubated for 2–3 hours at room temperature with continuous bubbling. This incubation was performed in ACSF with or without the desired compounds, always in the presence of a synaptic blocker cocktail (25 µM APV, 10 µM CNQX, 25 µM picrotoxin, 0.3 µM CGP 55845, and 1 µM DHβE) to eliminate glutamate-, GABA-, and acetylcholine-mediated transmission. For recording, slices were transferred to a recording chamber and continuously perfused at 34 °C with the synaptic blocker-containing ACSF (either with or without the test compounds).

To record evoked dopamine (DA) release, a carbon-fiber microelectrode (5 μm in diameter) was inserted into the dorsolateral striatum, and a bipolar stimulation electrode (WPI) was placed on the tissue surface approximately 50–100 μm away from the carbon fiber. FSCV data acquisition and analysis were conducted using Igor Pro 6 (Wavemetrics, Lake Oswego, OR) equipped with a custom-made FSCV macro (kindly provided by Dr. Eugene Mosharov). Background-subtracted triangular voltage waveforms (ramping from -450 mV to +800 mV at a scan rate of 294 V/s vs. an Ag/AgCl reference electrode) were applied to the microelectrode every 100 ms. Evoked DA peak currents were recorded using an Axopatch 200B amplifier (Molecular Devices) and digitized via an InstruTECH ITC-18 interface at a sampling rate of 10 kHz. To quantify the neurochemical responses, all carbon-fiber electrodes were individually calibrated with a 1 µM DA standard solution (Sigma-Aldrich) for each slice at the end of the session.

### Cloning and expression of full length SV2C, BRIL-SV2CΔ127, and SV2AΔ64

All genes were cloned using standard PCR primers and procedures. Briefly, full length SV2C was cloned into pEG-BacMam, containing an N-terminal 10xHis-StrepII-StrepII-mVenus tag linked to the SV2C N-terminus by a 3C protease site. To clone SV2CΔ127 the first 127 residues of the SV2C N-terminus were removed and replaced with the BRIL coding sequence, and this construct was cloned into pEG-BacMam containing an N-terminal 10xHis-StrepII-StrepII-mVenus tag linked to the BRIL N-terminus by a 3C protease site. SV2AΔ64 was truncated at the 64^th^ N-terminal residue and cloned into pEG-BacMam containing a C-terminal mVenus-StrepII-StrepII-10xHis tag linked to the SV2A C-terminus by a 3C protease site. All constructs were transformed into DH10Bac cells and used to generate baculovirus by previously established methods. Viruses were raised to >10^8^ infectious particles per mL by passaging in Sf9 cells. High-titer viruses were added at a ratio of 10% total culture volume to transduce tSA-201 cells at a density of 2.5-3.0x10^6^ cells/mL. 18-22 hours post-transduction, 10 mM sodium butyrate was added to cultures to enhance protein expression. Cells were harvested by centrifugation 72 hours post-transduction and pellets were stored at -80 degrees C.

### Fluorescence detection size exclusion chromatography

Fluorescence detection size exclusion chromatography (FSEC) was performed on a Shimadzu HPLC instrument equipped with a refrigerated autosampler and an RF20-AXS fluorescence detector with multi-wavelength detection, attached to a Superose 6 5/150 column. For mVenus tagged constructs, excitation and emission wavelengths were 515 nm and 528 nm, respectively. For tryptophan fluorescence of untagged constructs, excitation and emission wavelengths were 280 nm and 325 nm, respectively.

### Membrane production

Cell pellets were resuspended in TBS150 (20 mM Tris, pH 8.0, 150 mM NaCl) with protease inhibitors (200 nM aprotinin, 2 uM leupeptin, and 2 ug/mL pepstatin). The resuspended cells were kept cool on ice with constant stirring and sonicated for a total of 15 minutes at 40% power on a Fisherbrand 505 500 watt sonicator, 2 seconds on, 4 seconds off. Large debris was cleared with a 20 minute spin at 12,000x g followed by ultracentrifugation at 180,000x g for 2 hours to isolate membranes. Membranes were resuspended in TBS150 and homogenized with 25-30 passes in a Dounce homogenizer. Homogenized membranes were stored at -80 C for future use.

### Scintillation proximity assays

Adherent tSA-201 cells were transiently transfected with mVenus-SV2B or mVenus-SV2C constructs using PolyJet (SignaGen) according to the manufacturer’s instructions. After 48 hours, cells were harvested, washed with TBS150, and solubilized for 1 hour in TBS150 10 mM MNG and 1 mM CHS. The solubilization mixture was clarified by ultracentrifugation for 30 minutes at 100,000x g.

Concentration of each fluorescently tagged protein in the whole cell lysate was determined by monitoring mVenus fluorescence with FSEC. Whole cell lysates were diluted to a concentration of 5-10 nM SV2B/SV2C and incubated with ^3^H-padsevonil and 0.5 mg/mL of Cu-YSi scintillant beads (Revvity) in TBS150 containing 0.2 mM MNG. Nonspecific background counts were determined by measuring samples containing ^3^H-PSL and 100 uM unlabeled padsevonil. Samples were measured in triplicate in 96-well plates using a MicroBeta 2 scintillation counter and data was analyzed in GraphPad Prism 10 using the one-site specific binding option.

### Membrane filter assays

For binding experiments on single point SV2C mutants, tSA201 cells were transiently transfected with polyethyleneimine 40K using 1 ug of plasmid per mL of culture at a ratio of 3:1 PEI:DNA. Cells were harvested after 72 hours. Membranes, prepared as described above, were incubated with 5 nM ^3^H-PSL and varying concentrations of competitor ligands for 1 hour on ice in TBS150. 100 uL of 2.5% polyethyleneimine in TBS150 was passed through each filter well of a 96-well glass fiber C filter plate (Revvity) using a vacuum manifold. After incubation, membranes were passed through the filters and washed 4 times with 100 uL of TBS150. Nonspecific background counts were determined by measuring samples containing 5 nM ^3^H-PSL and 100 uM unlabeled PSL. Plates were dried overnight. The next day, 30 uL of scintillant was added to each filter, and the plate was counted in a MicroBeta 2 scintillation counter. Data was analyzed in GraphPad Prism 10 using the K_i_ fitting option.

SV2AΔ64-mVenus wild-type or SV2A Δ64-C297A-mVenus mutant plasmids were transfected into tsA201 cells as described above. Protein expression of each construct was analyzed by FSEC, and membranes containing equal amounts of proteins were used in the filter binding assays. We prepared a working radioligand stock of ^3^H-PRM by diluting concentrated ^3^H-PRM (66.67 µM, 15 Ci/mmol) using unlabeled PRM to a final concentration of 4 nM and 56 nM, respectively. For saturation-binding studies, SV2AΔ64-mVenus and SV2A Δ64-C297A-mVenus enriched membranes were incubated for 1 hour in presence of twofold serially diluted ^3^H-PRM (60nM to 0.46 nM) in triplicate. Specific binding of ^3^H-PRM to SV2AΔ64-mVenus or C297A mutant membranes was calculated by subtracting nonspecific binding from the total binding.

Nonspecific binding was measured by residual binding observed in the presence of 100 uM PSL. The K_d_ values were calculated by GraphPad Prism 10 software using nonlinear fitting for one site-specific binding.

### Wild type SV2C and BRIL-SV2CΔ127 purification

Full length SV2C membranes were solubilized in TBS150 containing 10 mM lauryl maltose neopentyl glycol (MNG), 1 mM cholesteryl hemisuccinate (CHS), 1 uM padsevonil (PSL), and protease inhibitors for 1.5 hours at 4 degrees C by stirring. Solubilization mixture was clarified by ultracentrifugation at 180,000x g for 1 hour. Clarified solubilization mixture was batch bound to Strep-Tactin resin for 1 hour at 4 degrees C and passed through a manual gravity-flow column. Bound SV2C was washed with 10 column volumes of TBS150 containing 0.2 mM glycodiosgenin (GDN) and 1 uM PSL. Protein was eluted from the column using TBS150 containing 0.2 mM GDN, 1 uM PSL, and 5 mM desthiobiotin.

Purification tags were removed from SV2C by overnight incubation with 3C protease at 4 degrees C. After tag removal, protein was concentrated and size exclusion chromatography was performed using a Superose 6 10/300 column. Fractions were checked for homogeneity using FSEC, monitoring Trp fluorescence. Peak fractions were concentrated to 8 mg/mL and 1 mM PSL was added just before freezing. For purification of BRIL-SV2CΔ127, the same procedure was used with the following modifications: protein was solubilized in TBS150 containing 20 mM dodecyl maltoside and 1 mM CHS, protein was purified apo and ligand was added just prior to grid freezing, and BAG2 anti-BRIL Fab was added at a 3 to 1 molar ratio of Fab:SV2C before the size exclusion step. The BAG2 anti-BRIL Fab was purified according to established methods^64^.

### SV2A purification

SV2AΔ64 purification was carried out as described previously^42,43^. In brief, cell membranes were resuspended in TBS_150_ and incubated with 5 µM of PRM for 30 min on ice, followed by solubilization using 10 mM lauryl maltose neopentyl glycol (LMNG) and 1 mM cholesteryl hemisuccinate (CHS) at 4 °C for 90 min. The insolubilized fraction was removed by ultracentrifugation at 40,000 rpm for 60 min at 4 °C, and the cleared supernatant was loaded onto a gravity-flow column containing GFP-nanobody conjugated sepharose resin. The resin was washed with 20 column volumes of TBS_150_ containing 0.020 mM CHS, 0.2 mM glyco-diosgenin (GDN), and 2 µM PRM. The resin bound with SV2AΔ64 was incubated overnight with 3C protease to remove mVenus, TwinStrep, and 10-His-tags. The next day, the flow-through containing SV2AΔ64 was mixed with 4-to-5-fold molar excess of 8667-Nb, followed by concentration to 500 µl using a 100 kDa concentrator (Amicon). The SV2A-PRM-8667-Nb complex was injected onto a Superose 6 10/300 column (Cytiva) in TBS_150_ containing 0.020 mM CHS, 0.2 mM GDN, and 2 µM of PRM. Individual SEC fractions of the homogenous peak corresponding to monomeric SV2AΔ64-PRM-8667-nb were further analyzed by FSEC. SEC fractions that were homogenous based on FSEC analysis were pooled and concentrated to 8.15 mg/ml using a 100 kDa concentrator before addition of 1 mM of PRM. The concentrated sample was ultracentrifuged at 60,000 g prior to cryo-EM grid preparation.

### Cryo-EM sample preparation and data collection

2 uL of SV2A or SV2C complex was applied to glow discharged Quantifoil grids (1.2/1.3 and 2/1 200 mesh, gold or copper). Grids were blotted on an FEI Vitrobot MK IV, blot force 4, blot time 4 at 100% humidity. SV2A-PRM, SV2C-PSL and SV2C-UCB-F grids were imaged on the University of Pittsburgh’s Titan Krios G3i at 300 kV, equipped with a Falcon 4i direct electron detector and a Selectris energy filter set to 10 eV slit width. These data sets were collected in counting mode with a calibrated pixel size of 0.719 Å per pixel and a defocus range of -0.5 to -2.5 um. The total dose for data collection was 50 e-/Å^2^. The apo SV2C dataset was collected at the Pacific Northwest Center for Cryo-EM on Titan Krios #1 at 300 kV using a Falcon 4i and a Selectris X (10 eV slit width) in counting mode with a calibrated pixel size of 0.7296 Å per pixel and a total exposure of 50 e-/Å^2^. Defocus was varied between –0.5 and –2.5 um.

### Cryo-EM data processing and model building

A total of 13,212 micrographs were collected for SV2A-PRM and 12,568 micrographs were selected for further data processing. We collected 46,200 micrographs for SV2C-PSL, 20,992 micrographs for SV2C-UCB-F, and 36,766 micrographs for apo-SV2C. CryoSPARC version 4.7.0 was used for initial processing. Micrographs were patch motion corrected and CTF estimated, followed by an initial round of blob picking. Blob picked particles were extracted at a box size of 384 for SV2A and 340 for SV2C, binned to box size 128 for initial processing steps. Blob picked particles were run through one round of 2D classification and a low-resolution ab-initio class was generated from selected 2D particles. Micrographs were denoised in CryoSPARC and the ab-initio model was used to generate templates for particle picking. After template picking on de-noised micrographs, 14,260,676 particles were extracted for SV2C-PSL, 3,076,623 particles were extracted for SV2C-UCBF, 3,488,233 particles were extracted for apo-SV2C, and 3,152,459 particles were extracted for SV2A-PRM, respectively. Particles were subjected to 1 round of heterogenous refinement against one good ab-initio volume with strong SV2 features and several decoy volumes of empty micelles and bad particles to remove junk. Hetero-refined particles were then used for an ab-initio reconstruction using the HR-HAIR method as described in Kim et al^76^. Outputs from this job that displayed strong transmembrane domain features and good ligand density were re-extracted at full box size and run through one round of non-uniform refinement. Non-uniform refined volumes were used to run more rounds of heterogenous refinement against a high quality SV2 volume and several decoy volumes as described above.

Once the particle stack was sufficiently clean, particles were exported to RELION 5.0 for Bayesian polishing. Polished particles were subjected to several rounds of RELION 3D classification, followed by importing back into CryoSPARC for local refinement, CTF refinement, and sharpening. This produced 26,282 particles refining to 3.53 Å for SV2A-PRM, 70,940 particles refining to 3.56 Å for SV2C-PSL, 40,132 particles refining to 3.29 Å for SV2C-UCB-F, and 47,041 particles refining to 3.50 Å for apo-SV2C. For model building of SV2A, a previously published PDB, 8UO9 was fit into the density and used as a starting model. Model building was done with Coot and iteratively real space refined in Phenix (version 1.20.1). For model building of SV2C, the AlphaFold model AF-Q496J9-F1 was fit into the density and used as a starting model for manual fitting in Coot. The fit model was run through one round of RosettaCM followed by manual adjustments in Coot and ISOLDE, with iterative real space refinement performed in Phenix version 1.20.1^77–79^.

## Acknowledgements

This work was supported by SPARK-NS, NIEHS R01ES023839 (to G.W.M.) and a Parkinson’s Foundation Postdoctoral Research Fellowship PF-PRF-933478 (to M.B.), as well as a Young Investigator Grant from the Brain and Behavior Research Foundation (to J.A.C., grant 30153). We thank Ali Salahpour for valuable input on experiments and the manuscript. We thank James Conway and the Pittsburgh Center for Cryo-EM for their support. We thank Haejung Chung for assistance performing Western blot experiments. A portion of this research was supported by NIH grant R24GM154185 and performed at the Pacific Northwest Center for Cryo-EM (PNCC) with assistance from Janette Myers. Plasmids coding for BAG2 Fab and BRIL sequences were kindly provided by Cheng Zhang at the University of Pittsburgh.

---

| **Data collection and processing** | SV2C-PSL<br>PDB 36TI<br>EMDB 77840 | SV2C-UCB-F<br>PDB 36TJ<br>EMDB 77841 | Apo SV2C<br>PDB 36TK<br>EMDB 77842 | SV2A-PRM<br>PDB 36TM<br>EMDB 77845 |
| --- | --- | --- | --- | --- |
| Number of micrographs | 46,200 | 20,992 | 36,766 | 13,212 |
| Magnification | 194,000x | 194,000x | 191,885x | 194,000x |
| Voltage (kV) | 300 | 300 | 300 | 300 |
| Electron exposure (e-/Å<sup>2</sup>) | 50 | 50 | 50 | 50 |
| Defocus range (μM) | -0.5 to -2.5 | -0.5 to -2.5 | -0.5 to -2.5 | -0.5 to -2.5 |
| Pixel size (Å) | 0.719 | 0.719 | 0.7296 | 0.719 |
| Symmetry imposed | C1 | C1 | C1 | C1 |
| Initial particle images | 14,260,676 | 3,076,623 | 3,488,233 | 3,152,459 |
| Final particle images | 70,940 | 40,132 | 47,041 | 26,282 |
| Map resolution (Å)<br>FSC threshold | 3.56 | 3.29 | 3.5 | 3.53 |
| Map resolution range (Å) | 3.3-4.7 | 2.8-4.0 | 3.1-5.3 | 3.1-8.1 |
| **Refinement** |  |  |  |  |
| Initial model used | AF-Q496J9-F1 | AF-Q496J9-F1 | AF-Q496J9-F1 | PDB 8UO9 |
| Model resolution (Å)<br>FSC threshold | 4.15<br>0.5 | 3.79<br>0.5 | 3.67<br>0.5 | 4.05<br>0.5 |
| Model resolution range (Å) | 4.15-16.8 | 3.79-22.4 | 4.05-19.6 | 4.05-18.7 |
| Map sharpening *B* factor (Å<sup>2</sup>) | -145.6 | -93.8 | -121.3 | -92.6 |
| **Model composition** |  |  |  |  |
| Non-hydrogen atoms | 3235 | 3408 | 3346 | 3230 |
| Protein residues | 420 | 429 | 427 | 425 |
| Ligands | 1 | 1 | 0 | 2 |
| ***B* factors (Å<sup>2</sup>)** |  |  |  |  |
| Protein | 140.07 | 40.78 | 61.94 | 41.58 |
| Ligand | 127 | 39.74 | - | 39.28 |
| **RMS deviations** |  |  |  |  |
| Bond lengths (Å) | 0.005 | 0.003 | 0.003 | 0.003 |
| Bond angles (°) | 1.106 | 0.728 | 0.627 | 0.518 |
| **Validation** |  |  |  |  |
| Molprobtity score | 1.41 | 1.83 | 1.55 | 1.41 |
| Clash score | 2.82 | 9.95 | 5.36 | 7.52 |
| Poor rotamers (%) | 0 | 1.11 | 1.97 | 0 |
| **Ramachandran plot** |  |  |  |  |
| Favored (%) | 95.1 | 95.94 | 97.84 | 99.04 |
| Allowed (%) | 4.66 | 3.58 | 2.16 | 0.96 |
| Disallowed (%) | 0.25 | 0.48 | 0 | 0 |

