## Supplemental Materials for "Structural Determinants of Synaptic Vesicle Protein 2C Ligand Selectivity and Their Impact on Dopamine Release"

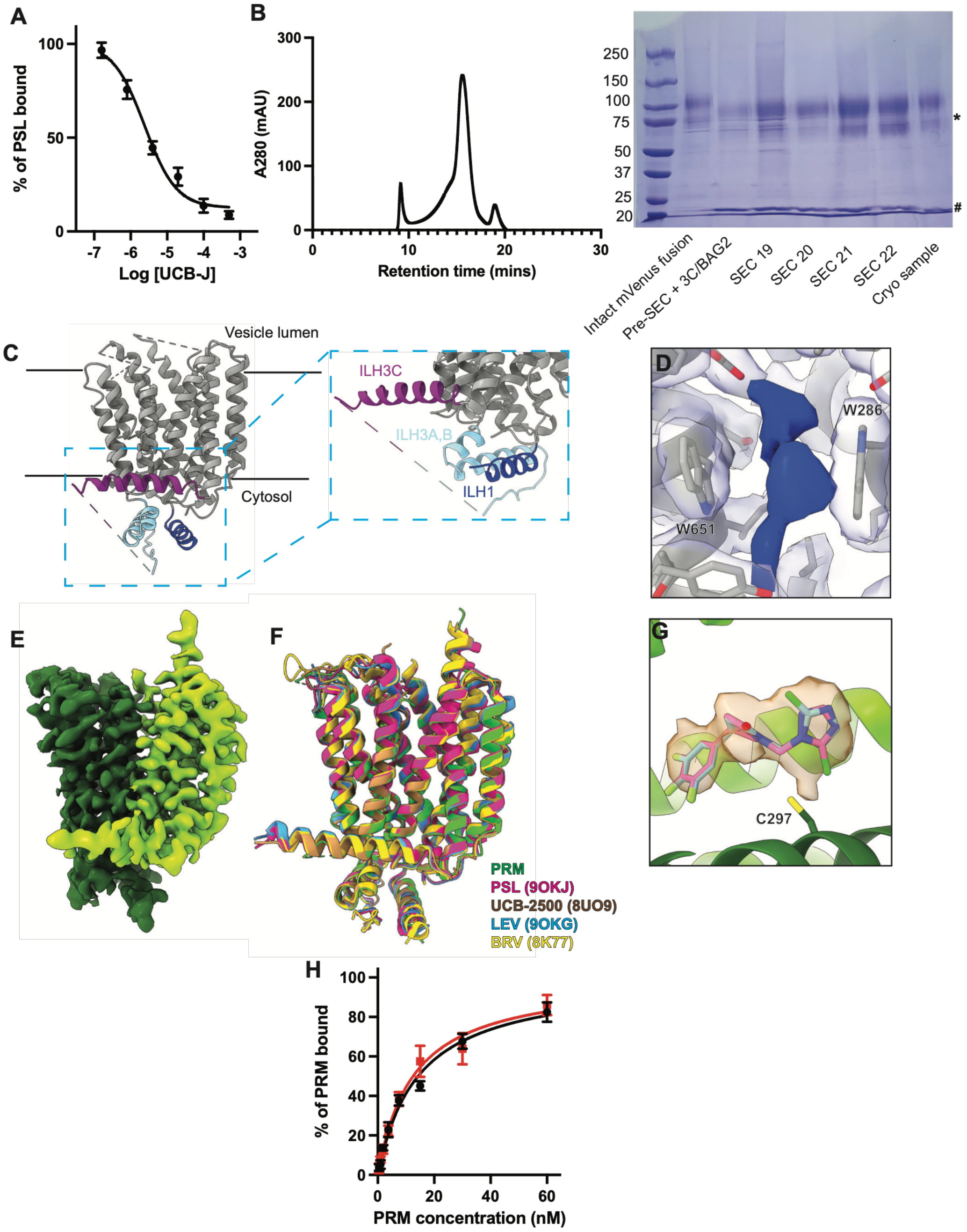


**Extended Data 1**

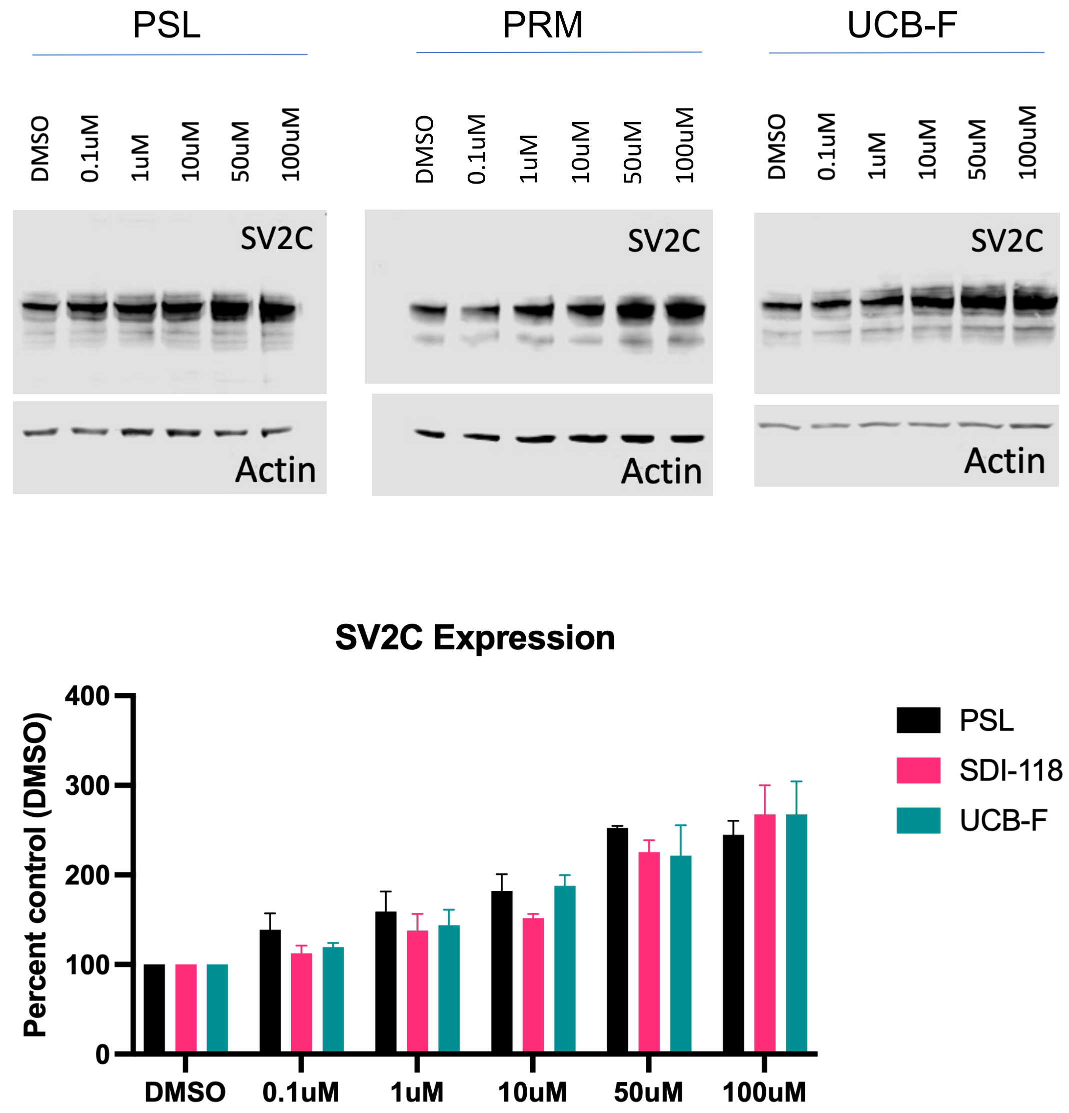


**Extended Data 2**

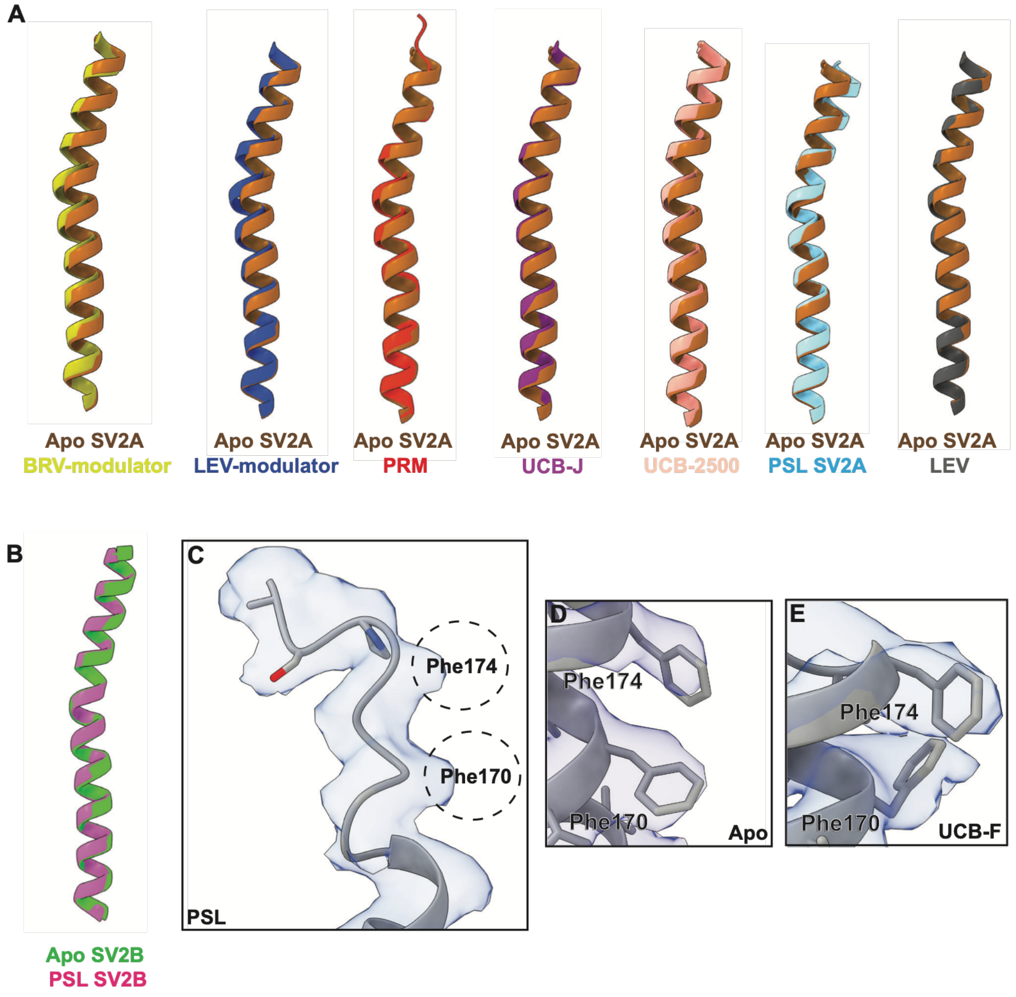


**Extended Data 3**

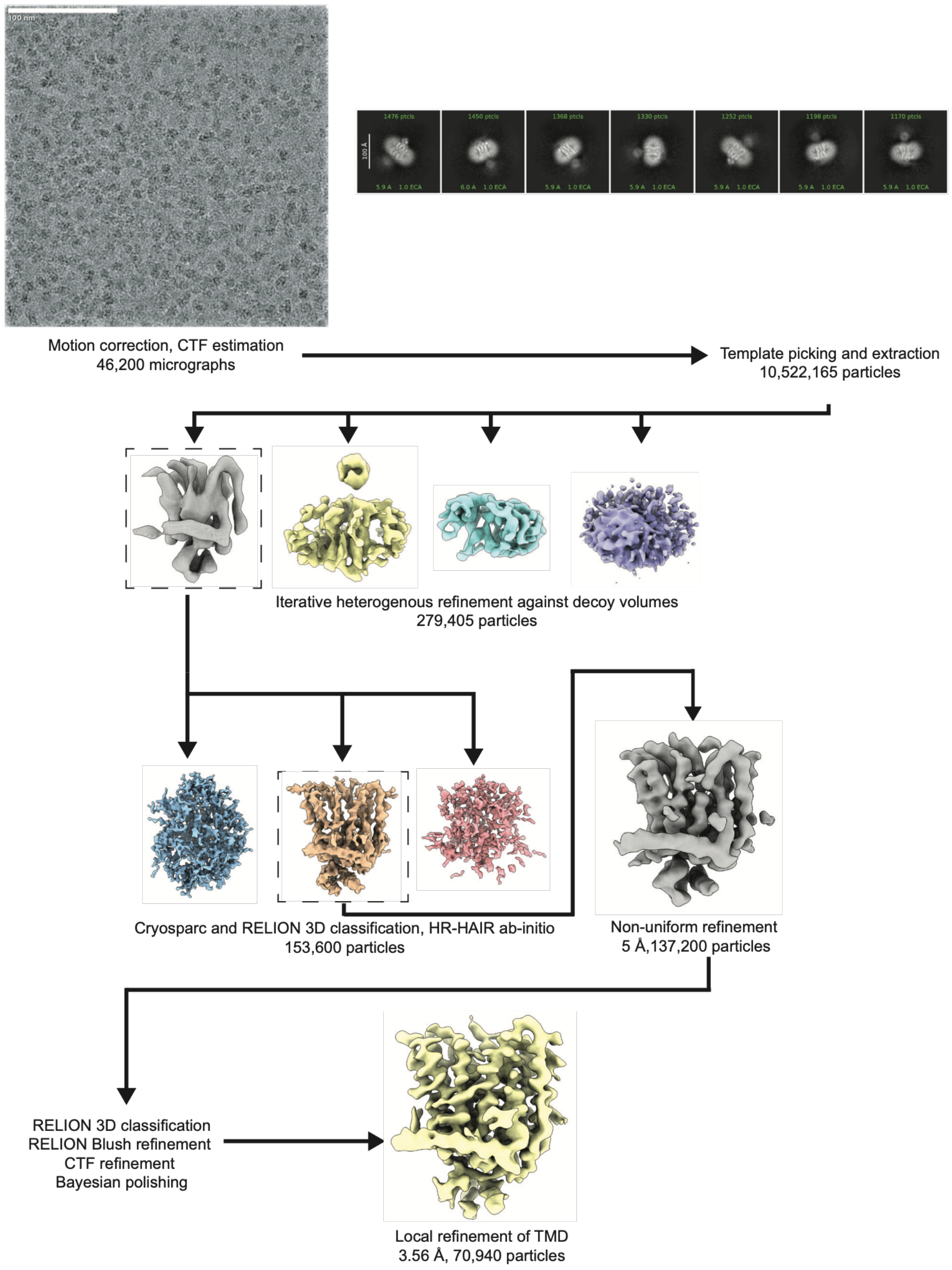


**Extended Data 4**

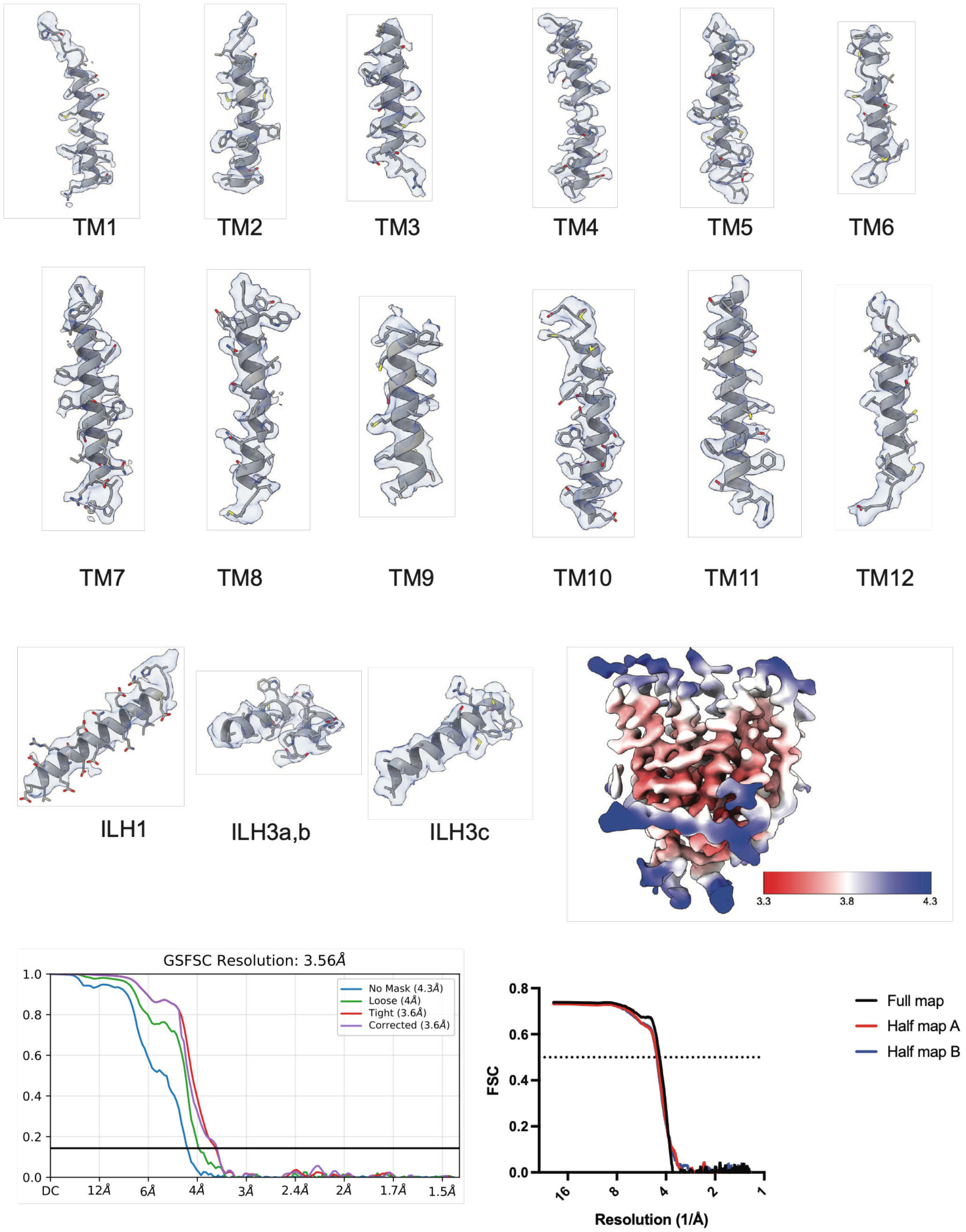


**Extended Data 5**

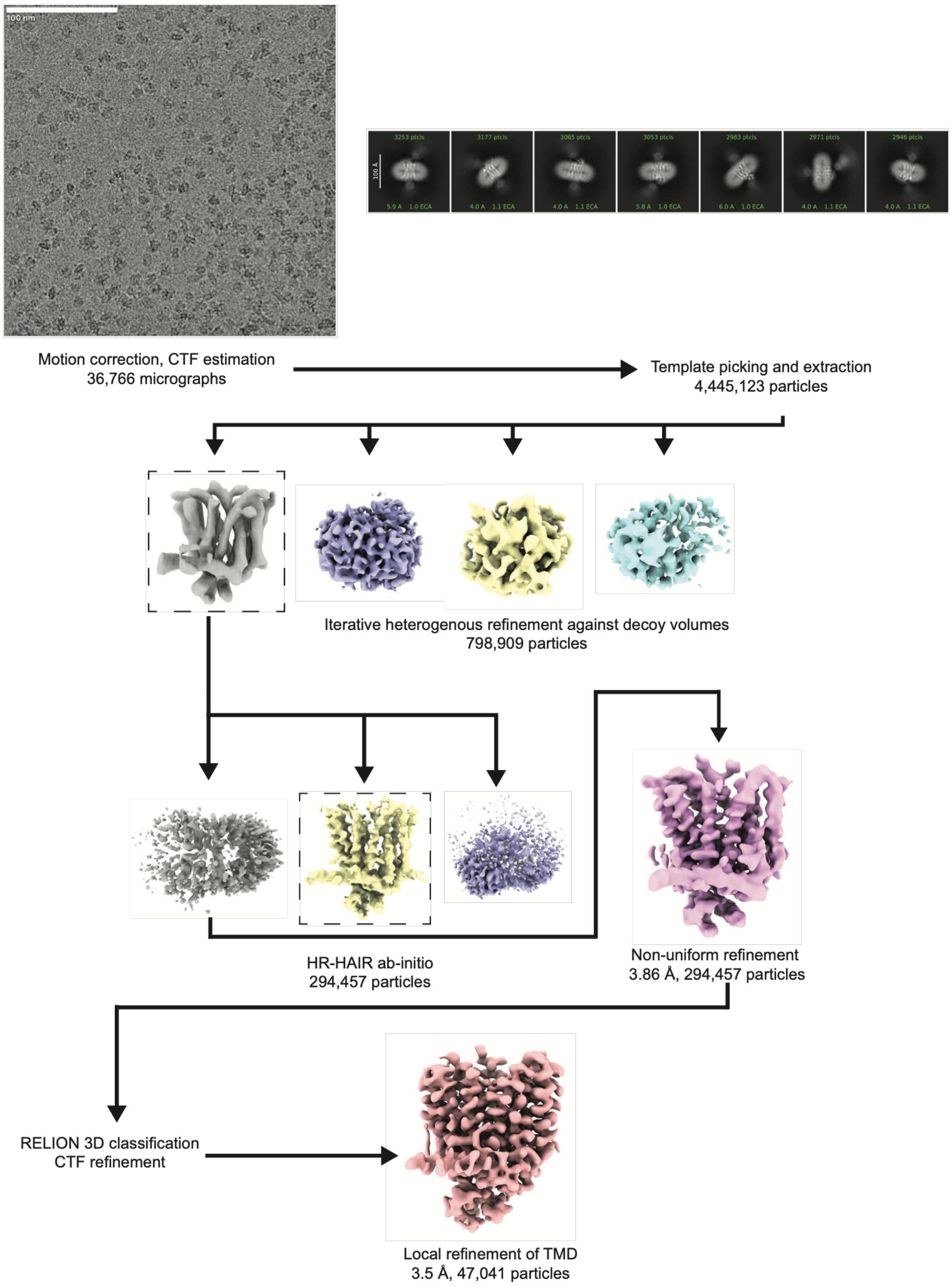


**Extended Data 6**

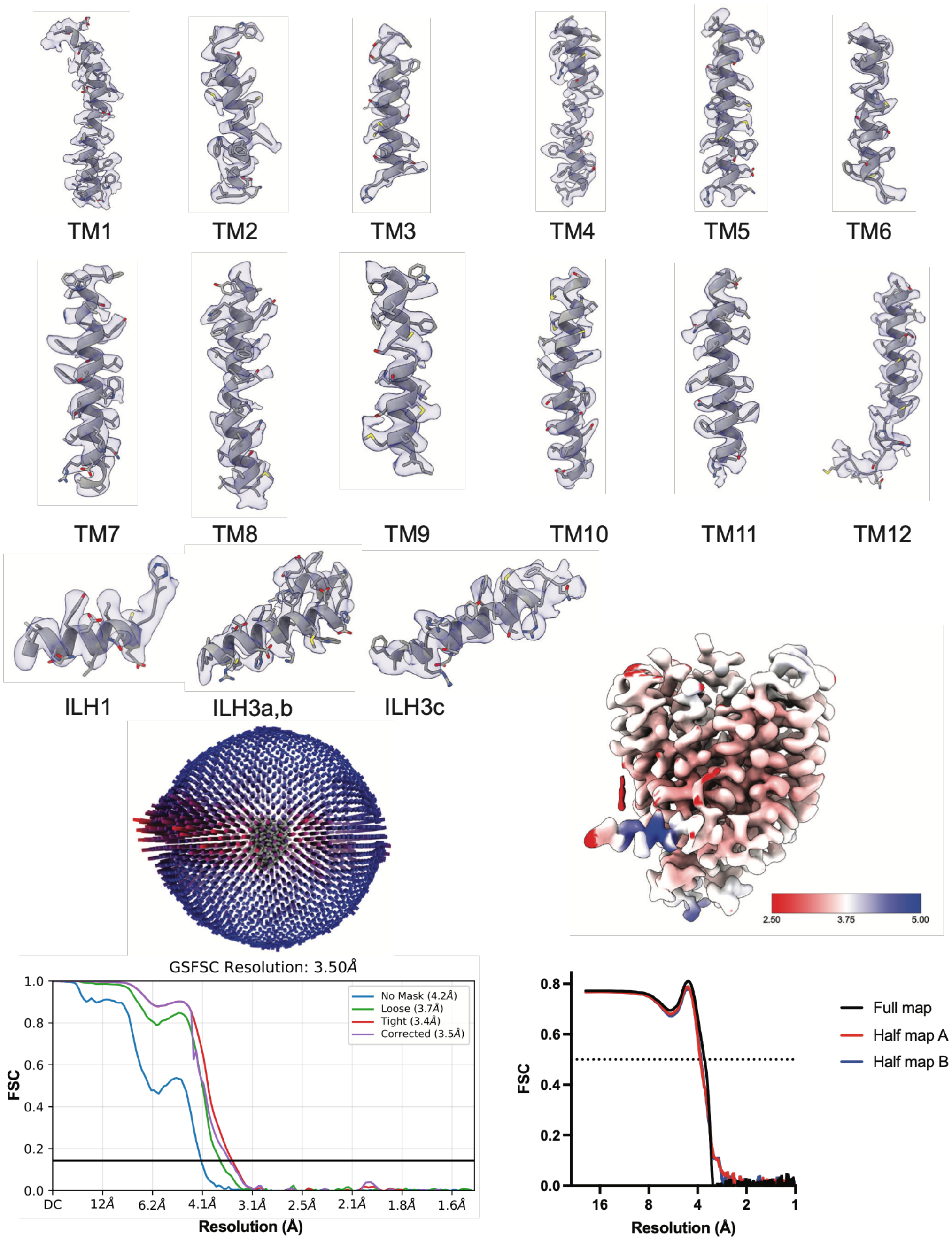


**Extended Data 7**

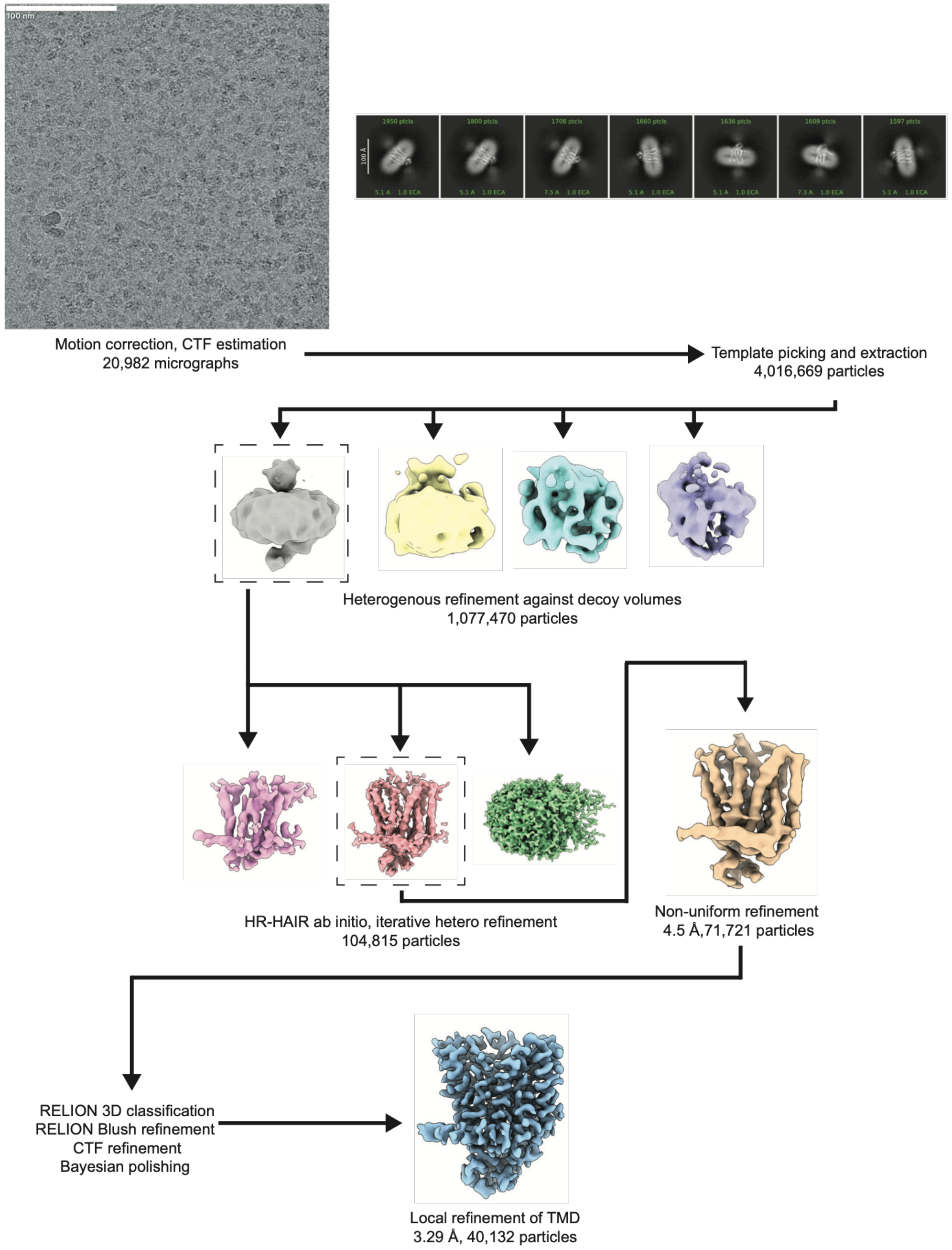


**Extended Data 8**

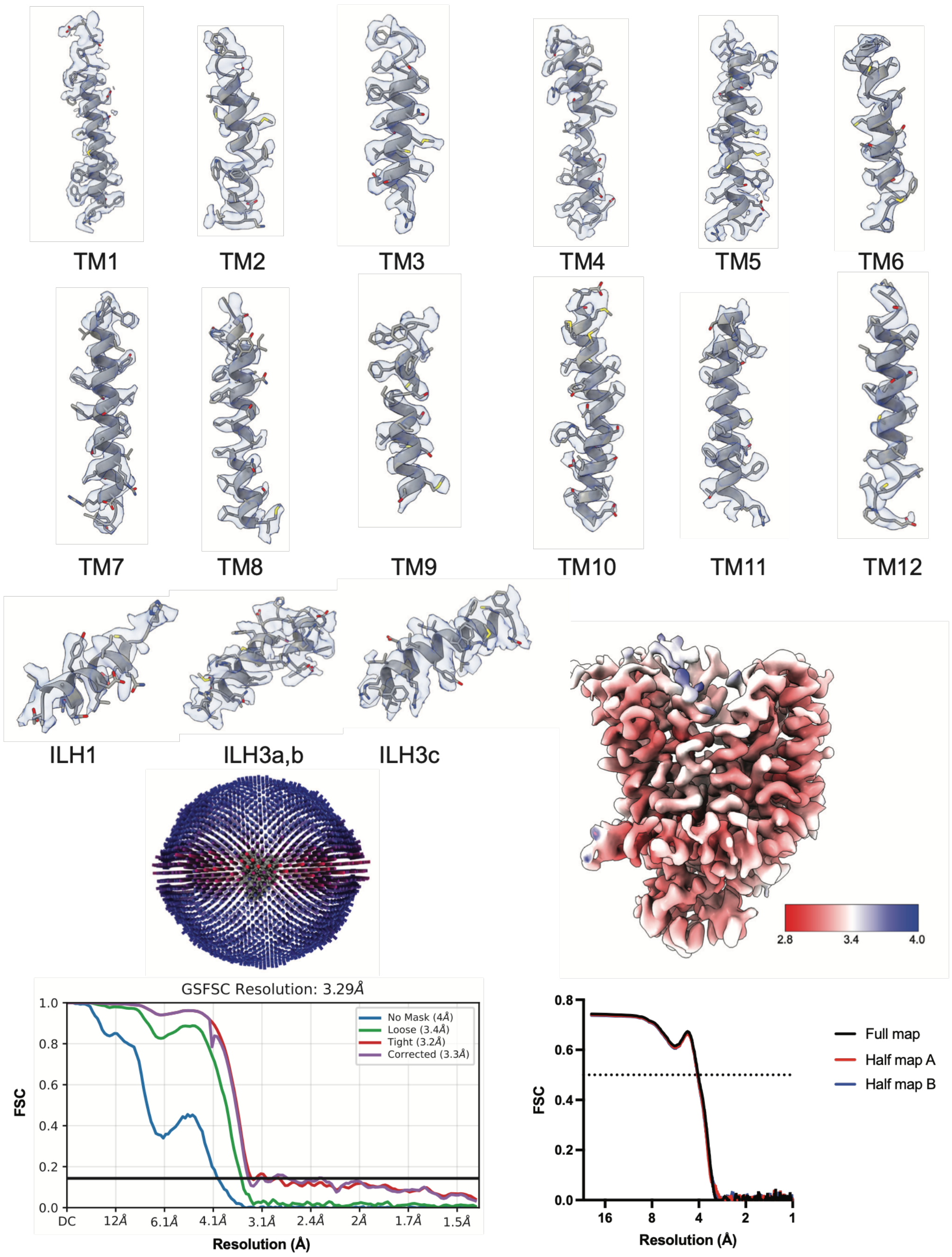


**Extended Data 9**

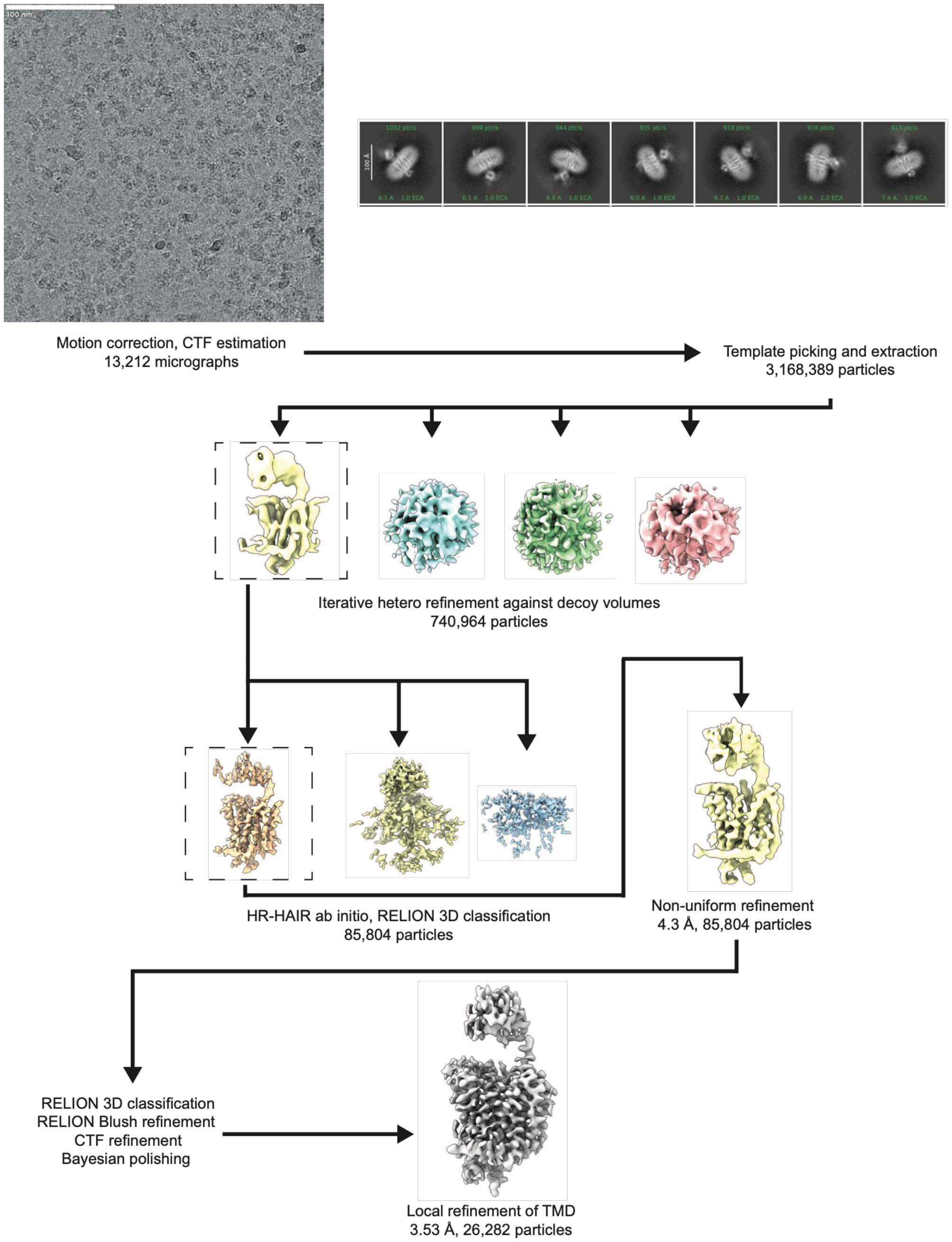


**Extended Data 10**

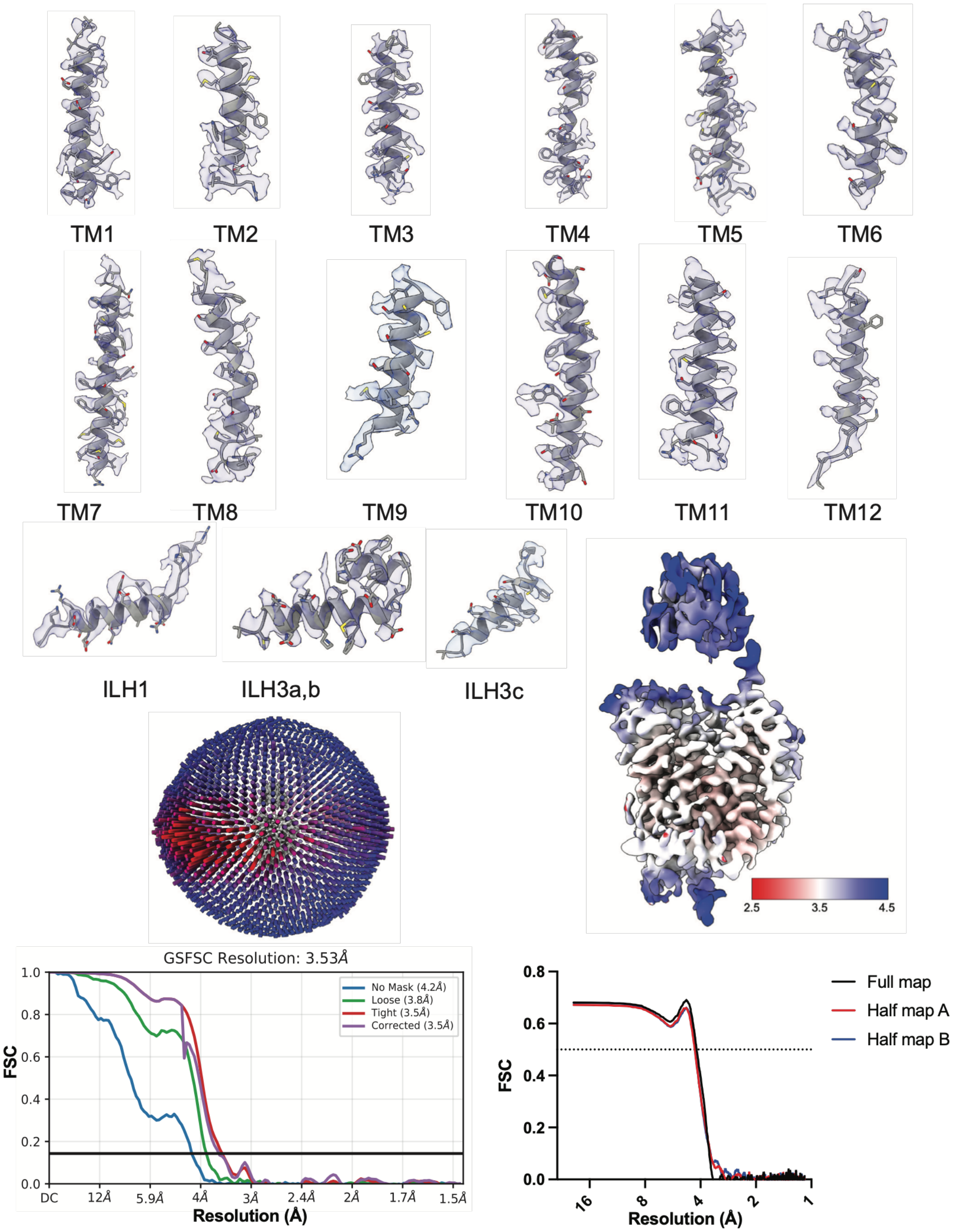


**Extended Data 11**
